# Evolutionarily conserved principles of ESCRT-III-mediated membrane remodelling revealed by a two-subunit Asgard archaeal system

**DOI:** 10.1101/2024.07.01.601590

**Authors:** Diorge P. Souza, Javier Espadas, Sami Chaaban, Edmund R. R. Moody, Tomoyuki Hatano, Mohan Balasubramanian, Tom A. Williams, Aurélien Roux, Buzz Baum

**Affiliations:** MRC Laboratory of Molecular Biology, Cambridge CB2 0QH, United Kingdom; Department of Biochemistry, University of Geneva, CH-1211 Geneva, Switzerland; School of Biological Sciences, University of Bristol, Bristol BS8 1TQ, United Kingdom; Centre for Mechanochemical Cell Biology, Division of Biomedical Sciences, Warwick Medical School, University of Warwick, Coventry CV4 7AL, United Kingdom

## Abstract

ESCRT-III proteins assemble into composite polymers that undergo stepwise changes in composition and structure to deform membranes across the tree of life. Here, using a phylogenetic analysis we demonstrate that the two ESCRT-III proteins present in our closest archaeal relatives are evolutionarily related to B-type and A-type eukaryotic paralogues, which initiate and execute membrane remodelling, respectively. This deep homology is reflected in ESCRT-III structure and function as demonstrated by the fact that ESCRT-IIIB assembles into parallel arrays on planar membranes to initiate membrane deformation, and is required to recruit ESCRT-IIIA to generate composite polymers. ESCRT-IIIA homopolymers can then remodel membranes into tubes, as a likely prelude to scission. Taken together, this analysis reveals a set of conserved principles governing ESCRT-III-dependent membrane remodelling that first emerged with the evolution of a two-component ESCRT-III system in the Asgard archaea, and which continue to underlie complex multi-component, ESCRT-III-dependent membrane remodelling in eukaryotes.

## Introduction

All cells and eukaryotic organelles are bounded by selectively permeable membranes that must be remodelled and repaired to maintain their organisation over time. To aid these processes, cells from across the tree of life employ a conserved family of polymer-forming ESCRT-III proteins that deform, cut and repair membranes.^1–3^ In eukaryotes, whose genomes tend to encode numerous ESCRT-III homologues, ESCRT-III proteins perform a host of important functions at different locations in the cell and in different contexts. For example, eukaryotic ESCRT-III proteins mediate cytokinetic abscission,^4,5^ viral budding,^6^ formation of endosomal multivesicular bodies for the degradation of ubiquitylated membrane proteins,^7^ plasma membrane repair,^8^ and nuclear envelope reformation following mitotic exit.^9,10^

The multitude of ESCRT-III homologues (12 subunits in humans and more than 25 in some plants^11^) are usually classified into 8 subfamilies: Vps46/Did2, Vps2, Vps24, Vps32/Snf7, Vps60, Vps20, CHMP7 and IST1 (CHMP1-7 and IST1 in animals, respectively)^7,12–17^. While subunits like CHMP4/Snf7 are known to be active in many cellular contexts,^3^ some have specific functions. Notably, CHMP7 is localised to the nuclear envelope where it aids reformation of the nuclear compartment at mitotic exit.^9,18^ At the same time, any one of these membrane remodelling events requires multiple ESCRT-III homologues acting in concert. As an example of this, the sequential recruitment of different ESCRT-III subunits is a common feature of endocytosis and other budding reactions, where it is thought to order and regulate membrane budding in time and space.^19^ A well characterised multi-step process of ESCRT-III-dependent membrane remodelling occurs during the formation of multivesicular bodies when CHMP6/Vps20 is recruited to the membrane, followed by CHMP4/Snf7, CHMP2A-CHMP3, and finally CHMP1B/IST1.^7,17,20–23^ In part because of the complexities of the eukaryotic ESCRT-III system, much remains to be understood about the general principles that govern the stepwise remodelling of membranes by ensembles of ESCRT-III proteins.

In this light, recent work has shown that homologues of eukaryotic ESCRT-III proteins perform a range of membrane remodelling functions in archaea, where they drive cell division ^24–26^, viral budding, and vesicle formation ^27–30^. Since archaea possess far fewer ESCRT-III homologues than most eukaryotes, they also provide useful simple models for studying ESCRT-III biology. Interestingly, the most stripped-down version of the ESCRT-III system is found in Asgard archaea, which are also the closest prokaryotic relatives of eukaryotes.^31–33^ Asgard archaea possess as few as two ESCRT-III homologues.^32–34^ While the difficulties in isolating and cultivating these organisms^35,36^ means that it is not yet possible to study this minimal system in cells, the two-component Asgard ESCRT-III system provides an excellent simple model for exploring the evolution,^34,37^ structure and function of ESCRT-III-dependent membrane remodelling.

Individual ESCRT-III subunits are small, consisting of 200-250 residues. These proteins have a relatively simple structure, with a set of alpha helices (α1-α5) that define the functional core of the protein, together with additional alpha-helical extensions at the N- and/or C-termini.^38,39^ These helices can be arranged in a so-called “closed” conformation, in which the alpha helices pack up against one another^40,41^ or in an “open” conformation.^42,43^ While there are instances in which polymers are formed by closed ^42,44^ or semi-closed subunits,^45^ in most cases polymers are constructed from subunits in the open conformation, which facilitates long range interactions between monomers within protofilaments. In these instances, adjacent subunits make numerous contacts with one another (mediated by helices α1 to α4), including a highly conserved interaction between subunit *i* (helix α5) and subunit *i+4* (α1-2 hairpin).^42,44,46^ In most cases, the formation of ESCRT-III polymers is aided by membrane binding, in part because ESCRT-III proteins physically associate with negatively charged lipids.^38,47^ Furthermore, since each ESCRT-III homopolymer binds preferentially to membranes of a defined curvature, it is thought that the sequential binding and polymerisation of different ESCRT-III subunits is able to progressively reshape flat membranes into tubes.^21,48,49^ Nevertheless, the specific mechanism by which this occurs remains poorly understood.

Here, to better understand the mechanism by which ESCRT-III proteins bind to and remodel membranes we have chosen to use phylogenetic, biochemical, biophysical and structural techniques to characterise the two-subunit ESCRT-III system identified in a member of the Asgard archaea, Heimdallarchaeota archaeon AB_125. Importantly, this is the simplest possible model with which to explore the mechanism of step-wise ESCRT-III dependent membrane remodelling. Our phylogenetic analysis shows that the two Asgard ESCRT-III proteins are related to two functionally distinct sub-groups of eukaryotic ESCRT-III proteins that act early and late in the membrane remodelling process. Using cryo-electron microscopy (cryo-EM) to solve the first high-resolution structure of a full-length early ESCRT-III protein (ESCRT-IIIB), our analysis also reveals how flexibility in the structure of the ESCRT-III monomer in its open confirmation accommodates it’s ability to form lattices that adapt to changes in membrane curvature. Consistent with this finding, using total internal reflection fluorescence (TIRF) microscopy we show that ESCRT-IIIB is able to bind to flat and slightly curved membranes, and can recruit ESCRT-IIIA, leading to the step-wise deformation of flat membranes into tubes. Taken together, our analysis shows how two Asgard ESCRT-III proteins can perform the same tasks as a large number of eukaryotic paralogues, and reveals common principles of ESCRT-III dependent membrane remodelling.

## Results

### Phylogenetic analysis of Asgard archaea and eukaryotic ESCRT-III proteins

Eukaryotes likely inherited ESCRT-III proteins from their archaeal ancestors. While many archaea possess multiple ESCRT-III paralogues, the genomes of the closest prokaryotic relatives of eukaryotes, Asgard archaea, encode as few as two.^1,32–34^ To better understand the evolutionary relationship between archaeal and eukaryotic homologues, we collected ESCRT-III sequences from Asgard and eukaryotes, including representatives of the complete set of eukaryotic ESCRT-III protein subfamilies, and performed a phylogenetic analysis (Figure 1A, S1). This revealed the existence of two main, maximally supported (100% bootstrap) clades of ESCRT-III proteins, each containing one Asgard ESCRT-III paralogue (ESCRT-IIIA and ESCRT-IIIB) and several eukaryotic subfamilies.^32,33^ The ESCRT-IIIA group (hereafter A-type) contains CHMP1/Vps46/Did2, CHMP2/Vps2, CHMP3/Vps24, and IST1, while the ESCRT-IIIB group (hereafter B-type) contains CHMP4/Vps32/Snf7, CHMP5/Vps60, CHMP6/Vps20, and CHMP7/CHM7.^1,32,50^ The topology of the tree, together with previous results,^1^ suggests a simple model of ESCRT-III evolution in which an ancestral gene was duplicated in the common ancestor of all Asgard archaea, giving rise to two specialised ESCRT-III proteins (Figure 1B). Over the course of several rounds of gene duplication and diversification during eukaryogenesis, these paralogues gave rise to the complete eukaryotic set (Figure 1B). Our analysis also suggests that a duplication of the ancestral ESCRT-IIIB gene resulted in the early versions of CHMP4/5 and CHMP6/7, which duplicated again to give rise to CHMP4/5/6/7 that diverged in modern eukaryotes (Figure 1B).

**Figure 1.**
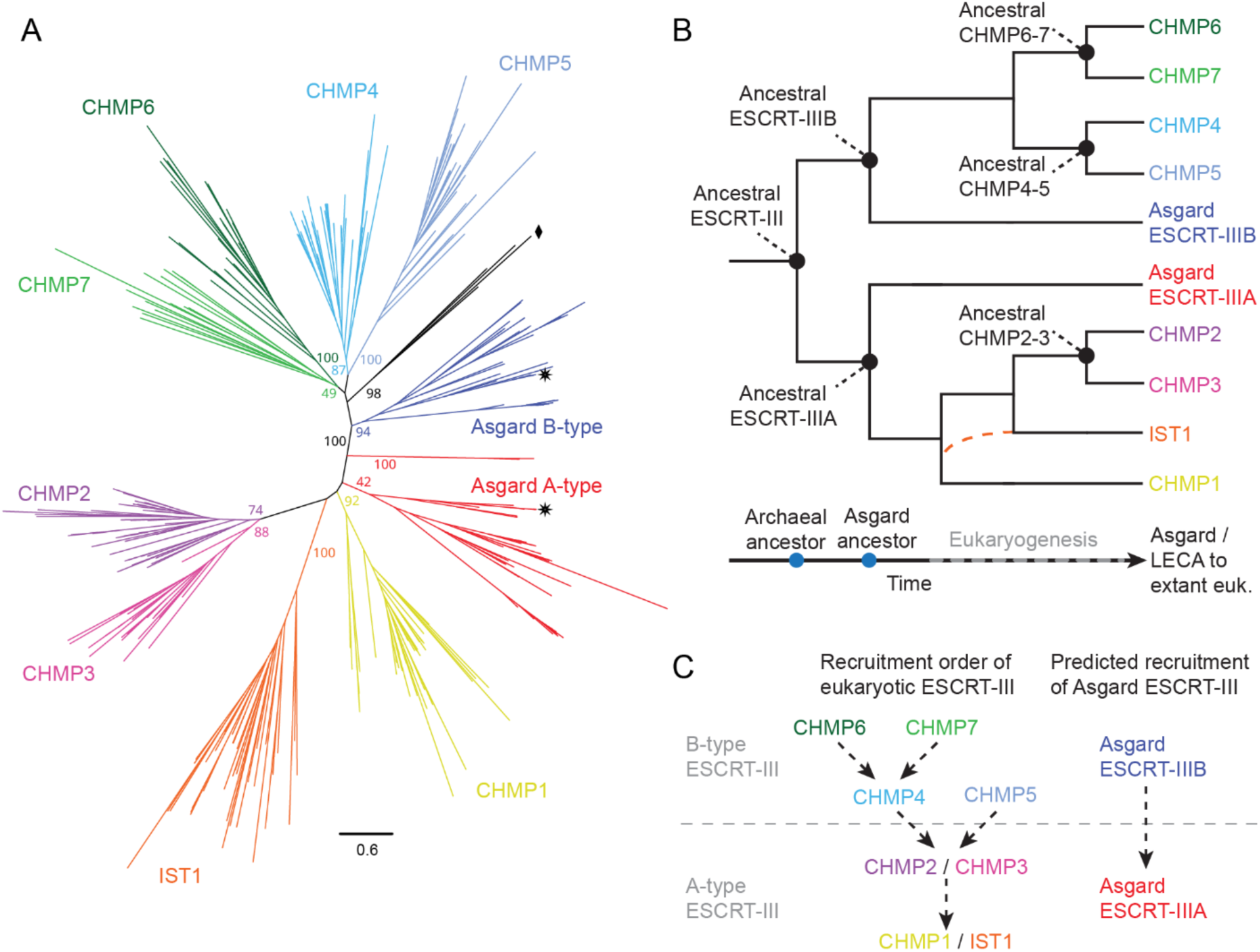
Evolution of the Asgard archaeal/eukaryotic ESCRT-III superfamily. A) Maximum likelihood phylogenetic tree of ESCRT-III proteins in eukaryotes and Asgard archaea. The A- and B-type proteins of Asgard archaea are related to eukaryotic CHMP1-3/IST1 and CHMP4-7, respectively. The A- and B-type sequences from Heimdallarchaeota archaeon AB_125, investigated experimentally in this study, are labelled with black stars. A small set of eukaryotic sequences lie outside of the main eukaryotic ESCRT-IIIB branch and are labelled with a black diamond (see Methods), implying the existence of a distinct group of ESCRT-III proteins. Support values are IQTREE2 ultrafast bootstrap supports.^79^ The scale bar represents the number of amino acid substitutions per site. A complete tree can be found in Figure S1. B) Proposed evolution of the eukaryotic ESCRT-III subfamilies based on the phylogeny shown in (A). The tree topology is consistent with an ancestral gene duplication in the common archaeal ancestor, followed by several rounds of gene duplication during eukaryogenesis, giving rise to all CHMP1-7/IST1 paralogues in the eukaryotic lineage prior to the last eukaryotic common ancestor (LECA). A proposal of when each ancestral ESCRT-III form appeared or existed is presented at the bottom. The position of IST1 in the eukaryotic A-type branch is not certain and could have followed pathways indicated by black or orange lines. euk.: eukaryotes. (C) The recruitment order of eukaryotic ESCRT-III proteins to membranes is related to the evolutionary history of its subfamilies. Taking into account the phylogeny (A-B) and the described recruitment order of their eukaryotic homologues (C, left), this analysis predicts that Asgard ESCRT-IIIB will be recruited to flat membranes before ESCRT-IIIA (C, right), which when recruited will induce membrane curvature.

Strikingly, an inspection of the subfamilies within the phylogenetic tree suggests that the ESCRT-III proteins within each of the two sister clades share functional and structural roles with one another. Proteins found in the ESCRT-IIIB family are typically recruited earlier in the membrane-remodelling pathway (CHMP4/5/6/7), while those in the ESCRT-IIIA family are recruited later (CHMP1/2/3/IST1) (Figure 1C). This suggests that the two ESCRT-III homologues found in Asgard archaea may share this recruitment order: one acting “before” (ESCRT-IIIB) and one acting “after” (ESCRT-IIIA) (Figure 1C). To test if this recruitment order predates eukaryogenesis and how only two Asgard ESCRT-III paralogues can perform the work of their eukaryotic counterparts, we investigated the structural and biophysical properties of ESCRT-III proteins encoded by one of our closest Asgard relatives, Heimdallarchaeota archaeon AB_125.^31,33^

### High-resolution structure of the Asgard ESCRT-IIIB protofilament

While much is known about the structure of A-type ESCRT-III subunits and protofilaments, the only known structures for B-type proteins are truncated versions of Snf7/CHMP4 solved by X-ray crystallography,^43,51^ and Snf7 protofilament spirals imaged by cryo-EM, which resolved the α1-α3 secondary structures.^52^ Therefore, to understand how B-type protofilaments are constructed, we began by purifying recombinant full-length ESCRT-IIIB from Asgard (Figure S2A), which was soluble and mostly monomeric (Figure S2B). However, after mixing it with large unilamellar vesicles (LUVs) containing 60 mol% of 1,2-dioleoyl-sn-glycero-3-phosphocholine (DOPC) and 40 mol% of the negatively charged lipid 1,2-dioleoyl-sn-glycero-3-phospho-L-serine (DOPS), long filaments were generated (Figure 2A). These filaments have a diameter of ∼12 nm and are reminiscent of those formed by Snf7/CHMP4/Shrub proteins.^53–55^ Filaments were also seen decorating and occasionally interacting with LUVs of various shapes, further suggesting that polymer nucleation is aided by the presence of membranes under these conditions (Figure 2A).

**Figure 2.**
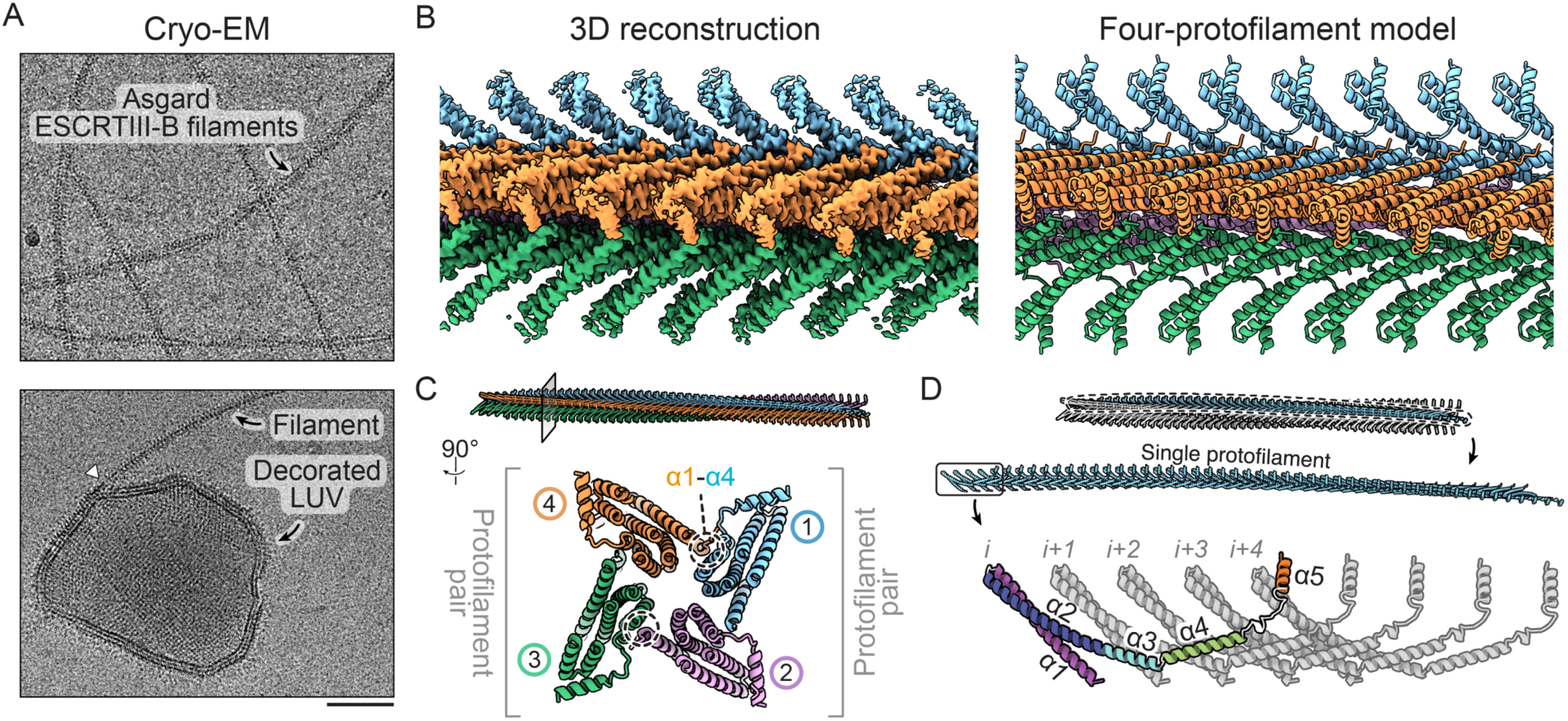
Asgard ESCRT-IIIB filament structure. A) Cryo-electron micrographs of Asgard ESCRT-IIIB bundles in the presence of LUVs. The bottom micrograph shows an example of a decorated LUV. An instance of a bundle connecting to a LUV is highlighted by the white triangle. (scale bar = 50 nm) B) Density map (left) and model fitting (right) of the Asgard ESCRT-IIIB bundle. The polymers are composed of four protofilaments (blue, pink, green and orange). C) Top view of a cross-section through the bundle, highlighting the two protofilament pairs and the inter-protofilament interactions through α1 and α4. D) Domain structure of the Asgard ESCRT-IIIB protofilament. Five subunits are shown (*i* to *i+4*). Secondary structure elements (helices α1-α5) of subunit *i* are highlighted.

The regularity of the filaments allowed us to use single particle analysis and helical reconstruction to obtain a density map at 2.9 Å resolution (Figure 2B, S3A-B, Table S1). Filament bundles were found to consist of four closely associated, parallel protofilaments arranged in two pairs with a shallow twist (∼1.5°) (Figure 2C, S3C). Each protofilament is built from monomers that interact through alpha helices α1-α5 (Figure 2D, S3C). Protofilaments are bound to their neighbours via electrostatic interactions linking helices α1 (N-terminus) and α4 (Figure S3D). Interestingly, the region of α4 mediating this interaction differs between adjacent pairs, resulting in a 1.5 nm offset (Figure S3C-D).

The monomers in these protofilaments adopt an open configuration (Figure S3C). Each monomer (*i*) makes extensive contacts with its neighbours at positions *i+1*, *i+2*, *i+3*, and *i+4* (Figure S3E), as described for subunits in A-type eukaryotic ESCRT-III protofilaments.^42,46^ This differs from the bacterial ESCRT-III homologues PspA and Vipp1, where interactions between subunits in a protofilament reach from *i* to *i+3,* even though the highly conserved interaction between α5 of subunit *i* and the hairpin α1/α2 are preserved.^1,56,57^ Therefore, this analysis of a B-type polymer shows that the underlying structural characteristics of ESCRT-III protofilaments are conserved from archaea to eukaryotes.

### The dynamics of Asgard ESCRT-IIIB binding to membranes

The requirement of LUVs for filament formation suggests that membranes trigger ESCRT-IIIB self-assembly. To directly observe this process, we fluorescently labelled ESCRT-IIIB and visualised its recruitment to supported lipid bilayers (SLBs) using TIRF microscopy (Figure 3A). While the presence of 20 mol% of negatively charged DOPS was sufficient to initiate the nucleation and growth of 0.5 ***μ***M yeast Snf7 (B-type) on SLBs,^55^ Asgard ESCRT-IIIB protein at the same concentration was not recruited to membranes containing ≤ 20 mol% DOPS (Figure 3A). Nevertheless, robust nucleation and growth of 0.5 μM ESCRT-IIIB was observed when SLBs contained ≥ 40 mol% DOPS (Figure 3A). On these DOPS-rich SLBs, Asgard ESCRT-IIIB initially formed a homogeneous layer within which small punctae appeared and grew until they completely covered the membrane. These data indicate that electrostatic interactions between ESCRT-IIIB and negatively charged head groups of lipids in membranes are important for nucleation, as they are for Snf7.^55^

**Figure 3.**
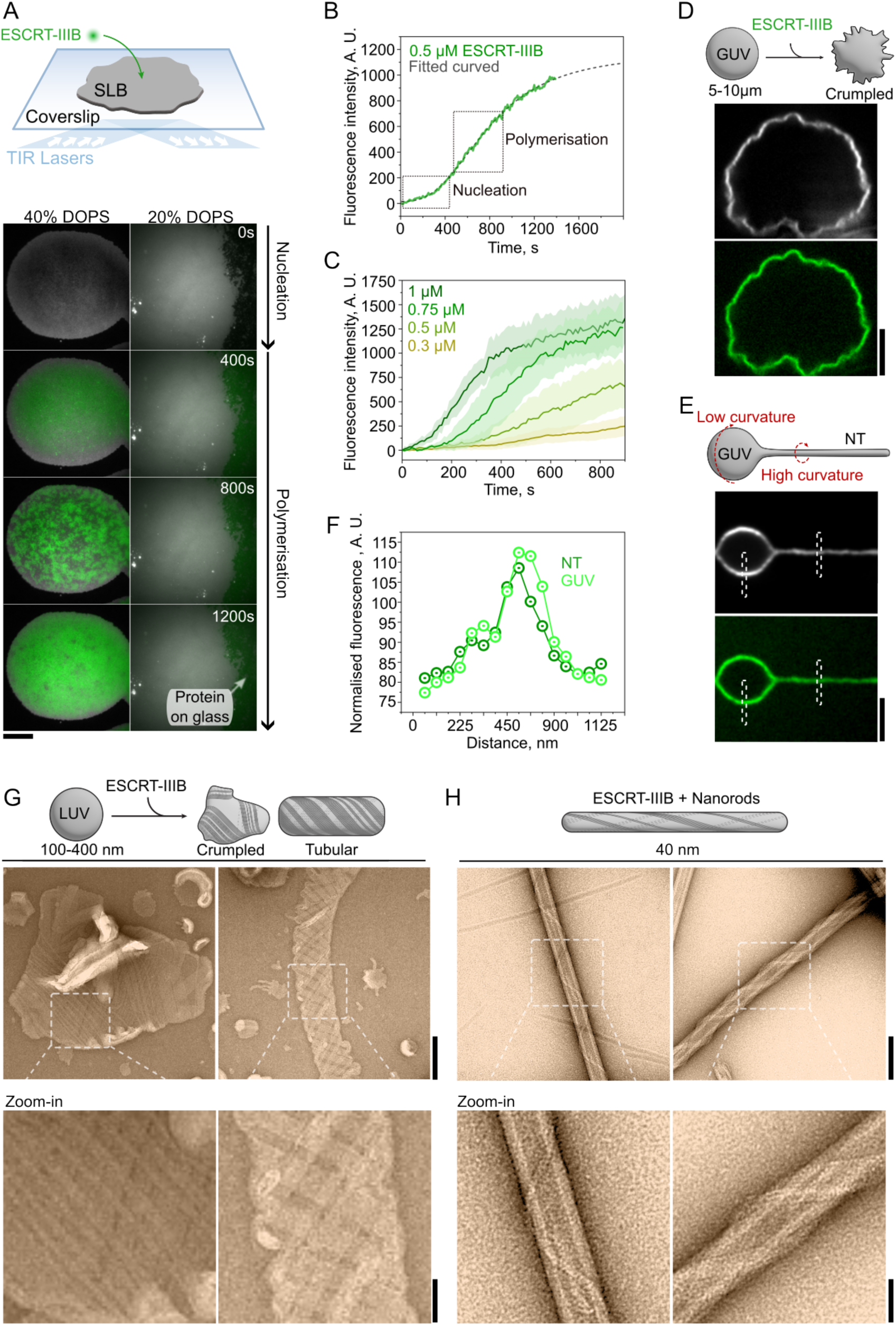
The interaction of Asgard ESCRT-IIIB with membranes. A) Schematic representation of the experimental set-up in which time-lapse fluorescence microscopy is used to follow ESCRT-IIIB (green) recruitment to flat SLBs (grey) over a period of 1200 seconds. Left and right columns show membranes containing 40 and 20 mol% of DOPS, respectively. ESCRT-IIIB binds and self-assembles on 40 mol% DOPS-containing membranes and fails to interact with 20 mol% DOPS-containing membranes, leading to its accumulation on the glass. Scale bar = 5 μm. B) Fluorescence signal time-profile of 0.5 μM ESCRT-IIIB over a SLB membrane. The green line shows Asgard ESCRT-IIIB mean fluorescence intensity (n=9 independent SLBs) measured on 40 mol% DOPS membranes (as shown in panel A). Left and right dashed boxes indicate ESCRT-IIIB nucleation and polymerisation, respectively. Dashed grey line shows the fitting curve using the Hill function. C) Fluorescence intensity time-profiles of Asgard ESCRT-IIIB measured on 40 mol% DOPS SLBs at concentrations where ESCRT-IIIB self-assembles (0.3, 0.5, 0.75 and 1 μM). (Mean and SD are shown for each concentration; n = 9 independent SLBs in all conditions). D) Schematic showing ESCRT-IIIB addition to a spherical GUV and a representative fluorescence image of Asgard ESCRT-IIIB induced membrane deformations (crumpling) upon complete surface coverage is shown. Scale bar = 3 μm. E) Schematic showing the GUV with a pulled NT. A representative fluorescence image of Asgard ESCRT-IIIB binding to a GUV and nanotube is shown. Binding is independent of membrane curvature. White dashed boxes show regions where the fluorescence plot profile was obtained from membrane and ESCRT-IIIB, to measure preferential protein recruitment dependent on membrane curvature and relative to membrane signal. In (D) and (E), the protein and membrane fluorescence signals are shown in green and grey, respectively. Scale bar = 2 μm. F) Fluorescence plot profiles of Asgard ESCRT-IIIB on flat GUV and curved lipid nanotube normalised versus membrane fluorescence profiles obtained at the same regions of interest (n=5 pulled NTs from different GUVs). Representative areas are shown in (E). G) Representative negative stain EM micrographs of Asgard ESCRT-IIIB filaments wrapping around flat and tubular LUVs and (H) nanorods. Scale bars are 100 nm and 40 nm in the zoom-out and zoom-in images, respectively.

An analysis of membrane-bound ESCRT-IIIB revealed a sigmoidal increase in fluorescence intensity over time (Figure 3B). The shape of the curve suggests a two-step kinetic process^58,59^ with an initial lag phase that is likely to reflect rate-limiting nucleation, followed by a rapid growth phase (i.e. monomer addition), until a slow-down occurs as membrane coverage reaches saturation (Figure 3B). In this way, ESCRT-IIIB appears to polymerise on the membrane in a manner similar to other cytoskeletal polymers like actin.^58^ Consistent with this interpretation, the process was concentration-dependent (Figure 3C). Below 200 nM, only a small uniform fluorescent signal was observed on the membrane, indicative of monomer binding without nucleation (Figure S4A). The growth rate of the polymers was proportional to the monomer concentration (Figure 3B, S4B, S4C), indicating the absence of cooperativity in the polymerisation process. Membrane-bound ESCRT-IIIB fluorescence intensity increased with time until it reached a final steady-state fluorescence intensity that was similar regardless of the monomer concentration (Figure S4D, S4E), suggesting that it is surface accessibility rather than bulk depletion that limits further polymer assembly.

To understand how the self-assembly of Asgard ESCRT-IIIB affects membrane geometry, giant unilamellar vesicles (GUVs) were mixed with the protein at levels greater than the critical concentration required for self-assembly, resulting in complete decoration by the protein. Under these conditions, >30% of GUVs exhibited a “crumpled” appearance (Figure 3D), which was different to the appearance of GUVs coated with lower surface densities of ESCRT-IIIB (Figure S4F). This “crumpling” resembles the effect of adding Snf7 (B-type) to GUVs,^55^ and is likely due to the formation of a dense, flat and continuous protein coat on the membrane (Figure 2A), which compresses the GUV without imposing a specific curvature.

B-type proteins like Snf7 are preferentially recruited to relatively flat membranes,^55,60^ and tend to be excluded from the high curvature of pulled membrane nanotubes (NTs).^21,55,60^ To test whether Asgard ESCRT-IIIB acts in this way, we pulled membrane NTs from spherical GUVs and compared ESCRT-IIIB fluorescence intensity on different parts of the membrane (Figure 3E). Strikingly, while ESCRT-IIIB binds flat membranes like Snf7, it also binds to highly curved NTs (with a radius of curvature of > 1000 nm in the spherical GUV versus ∼50 nm in the pulled NTs). Moreover, when normalised relative to membrane intensity, levels of ESCRT-IIIB fluorescence appeared similar in both regions of the membrane (Figure 3F). These data suggest that Asgard ESCRT-IIIB polymers are able to adapt to a wide range of membrane curvatures.

To understand how ESCRT-IIIB binds membranes of higher curvature, we performed negative stain EM to visualise its association with LUVs (100-400 nm in diameter) and thin cylindrical galactocerebroside nanorods (∼40 nm in diameter) (Figure S4G-H).^61,62^ On LUVs, Asgard ESCRT-IIIB polymers formed large lattices with angular edges (Figure 3G), likely corresponding to the “crumpled” structures seen on the larger GUVs (Figure 3D). In addition, we observed instances in which ESCRT-IIIB filaments wrapped around wide, slightly constricted membrane tubes to form parallel, tilted arrays that resemble a mesh in the projected 2D micrographs (Figure 3G). Conversely, when Asgard ESCRT-IIIB was added to nanorods, filaments did not wrap around the membrane to form tight helices. Instead, filaments formed extended membrane-bound helices that contained variable numbers of protofilaments (Figure 3H), that lay nearly parallel to the tube axis, resembling those previously observed for a mix of B-type with A-type proteins (Snf7-Vps2-Vps24).^63^ These data confirm that, in contrast to its eukaryotic counterparts, the polymers formed by Heimdallarchaeota ESCRT-IIIB are flexible enough to accommodate a wide range of membrane curvatures^54,55^ (Figure S4I).

### Structure of Asgard ESCRT-IIIB membrane arrays

Next, we attempted to solve the structure of Asgard ESCRT-IIIB filament arrays bound to membranes (Figure S5A). Because the membrane signal made it difficult to accurately align the arrays, we devised a strategy to reduce the membrane signal in the micrographs (Figure S5A) using techniques inspired by studies on proteins decorating microtubules.^64,65^ This allowed us to generate 2D class averages where the arrays are more clearly defined (Figure 4A). Despite their structural variability, single particle analysis using these data yielded a map at 6.5 Å resolution, allowing us to confidently assign alpha helices (Figure 4B, S5B-C, Table S1).

**Figure 4.**
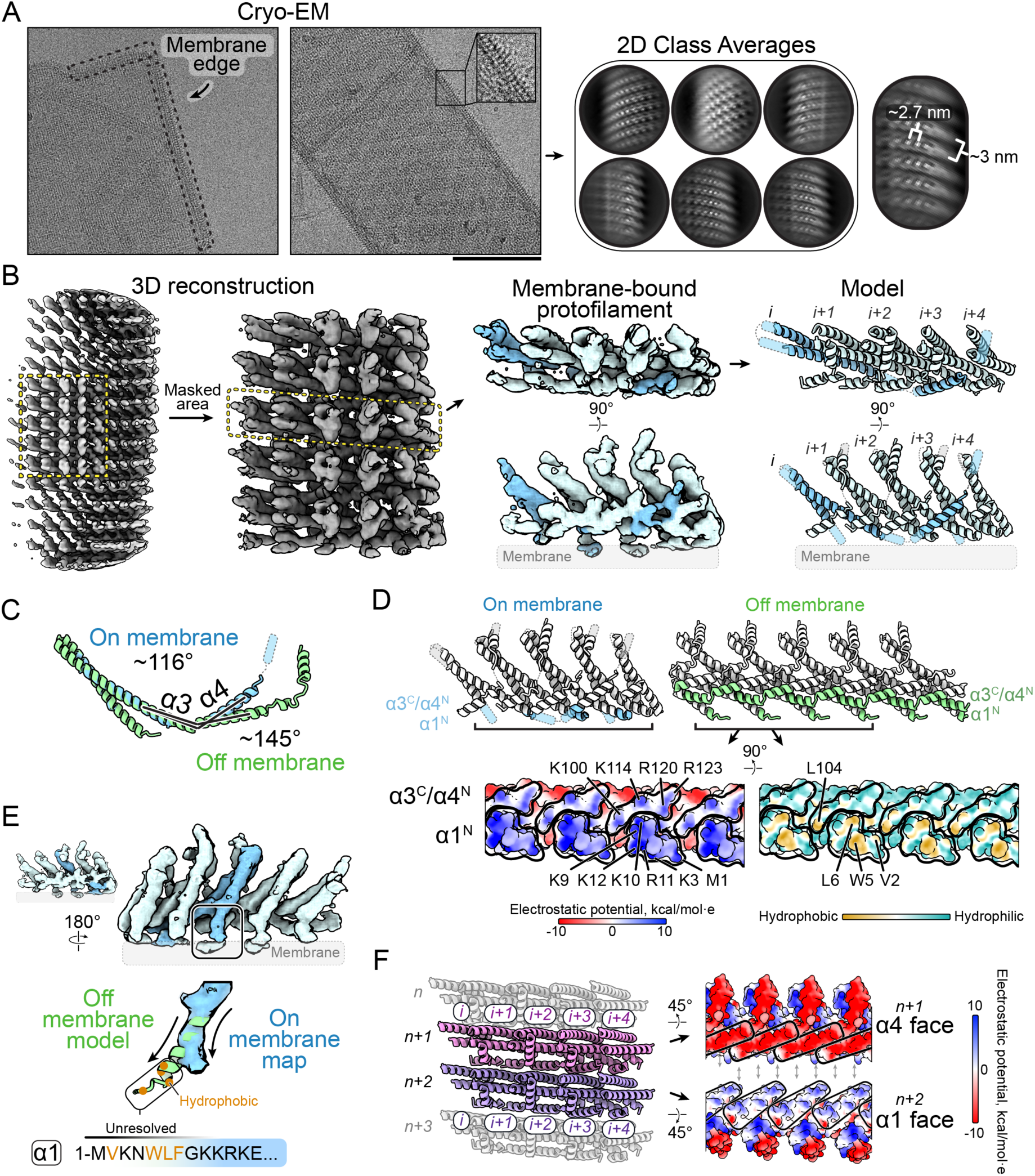
Structure of Asgard ESCRT-IIIB bound to membranes. A) Cryo-electron micrograph of Asgard ESCRT-IIIB decorating LUVs, with repeating spikes at the membrane edge (left; scale bar = 100 nm). Picked particles along the membrane edge are used for 2D classification (right). B) Density map of Asgard ESCRT-IIIB bound to membranes, showing the area masked for refinement (left). A single protofilament is highlighted along with the fit model (right). C) An overlay of the on-membrane and off- membrane models after aligning at α1-3, highlighting the angle between α3 and α4. D) Bottom view of the high-resolution Asgard ESCRT-IIIB off-membrane protofilament structure showing the charged/hydrophobic residues that would be exposed in the on-membrane model. E) A rotated view of the protofilament on membranes showing a deviating helix α1 and unresolved N-terminus, where a hydrophobic patch may extend into the membrane. F) Model of the polymeric arrangement of several Asgard ESCRT-IIIB protofilaments on LUVs (left). Lateral interactions between protofilaments are mediated by complementary charged surfaces in helices α1 (blue; positively charged surface) and α4 (red, negatively charged) (right).

A comparison with our off-membrane structure revealed a slightly smaller inter-subunit distance in the membrane-bound protofilament (∼3 nm versus 2.7 nm, respectively). This was accompanied by a change in the angle between helix α3 and α4 (off-membrane: 145°, on-membrane: 116°) (Figure 4C). This indicates that the loop between α3 and α4 acts as a hinge aiding the conformational flexibility of the protofilament - consistent with previous reports from eukaryotic and bacterial ESCRT-III polymers.^1,39,44^

The structure of membrane-bound Asgard ESCRT-IIIB also revealed that the protein binds membranes via the N-terminus of α1 and the loop connecting helices α3-α4 (Figure 4D). This interface includes an exposed region of continuous positive charge as well as hydrophobic residues (Figure 4D), consistent with the binding of filaments to negatively charged membranes (Figure 3A). By superimposing the high-resolution structure of α1 from our off-membrane model with our structure on membranes, we noticed that the hydrophobic N-terminus was positioned to insert into the lipid core of the membrane (Figure 4E). While this binding interface differs from the reported interaction site of CHMP1B (A-type) with membranes,^44,66^ it is similar to those of Snf7 (B-type)^52^ and CHMP2A-CHMP3 (A-type).^46^

The Asgard ESCRT-IIIB membrane interaction interface leaves the protofilament free to interact with adjacent protofilaments in the array. These lateral interactions are mediated by the positive charges of helix α1 and negative charges of α4 (Figure 4F). Interestingly, helical filaments formed by CHMP2A-CHMP3 use similar charge complementarity to mediate interactions between adjacent rungs.^46^ In addition, it was previously proposed that equivalent lateral interactions also connect composite polymers of B-type Snf7 and A-type Vps24.^67^ Although these interfaces differ from the ones that were previously proposed to mediate interactions between helical rungs of A-type CHMP1B,^42^ the data indicate that complementary charges located on extended surfaces are a general feature of B-type and A-type homo- and hetero-polymers - likely enabling protofilaments to flexibly associate with each other in ways that allow for filament sliding.^67^ Taken together, these data suggest that ESCRT-IIIB subunits and the protofilaments they form have an intrinsic flexibility that allows them to bind membranes of different curvatures.

### Asgard ESCRT-IIIA is recruited by ESCRT-IIIB

The observation that Asgard ESCRT-IIIB binds flat membranes is consistent with its role in initiating membrane remodelling, as suggested by the phylogenetic analysis. On the other hand, when we performed similar experiments with fluorescently labelled ESCRT-IIIA, it did not bind SLBs, even at high concentrations. Instead, after extended periods of time, the fluorescent protein was seen accumulating on the surrounding glass (Figure 5A, S2A).

**Figure 5.**
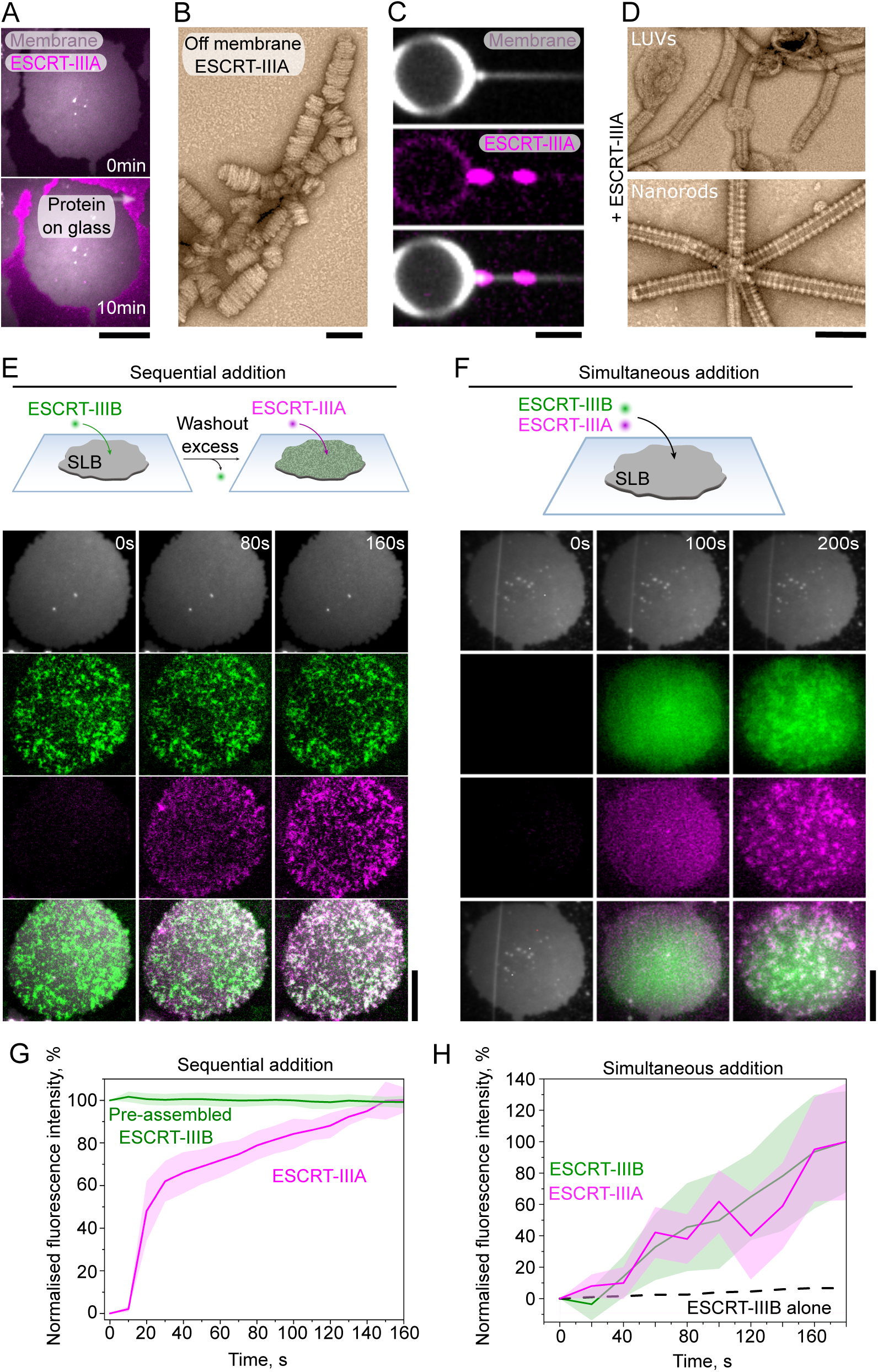
Characterisation of Asgard ESCRT-IIIA and its recruitment to flat membranes. A) Figure shows representative TIRF microscopy image which demonstrates that 1 μM Asgard ESCRT-IIIA is not recruited to 40 mol% DOPS SLB membranes after 10 min incubation. Scale bar = 10 μm. B) Representative negative stain electron micrographs of Asgard ESCRT-IIIA forming helical protein arrays when forming polymers without membranes. Scale bar = 100 nm. C) Fluorescence image shows 1 μM of fluorescently labelled ESCRT-IIIA forming clusters on the highly curved part of the membrane nanotube (pink; middle) that was pulled from a GUV (grey; top), after a 15 minute incubation and washing. Merge is shown at the bottom. Scale bar = 1 mm. D) Representative negative stain electron micrographs of Asgard ESCRT-IIIA on LUV and nanorod membranes. ESCRT-IIIA forms a helical tube on nanorods (bottom), whereas the protein self-assembly promotes LUVs tubulation (top). Scale bars are 110 nm. E) Schematic representation of the sequential addition experiment. Fluorescence time-lapse images were acquired using TIRF microscopy to show Asgard ESCRT-IIIA recruitment to the SLB membranes with pre-assembled ESCRT-IIIB patches. Scale bar = 5 μm. 1 μM of ESCRT-IIIB was added to form the patches and washed-out before complete membrane coverage, and then 1 μM of ESCRT-IIIA was added. F) Schematic representation of the simultaneous addition experiment. Fluorescence time-lapse images acquired using TIRF microscopy show how Asgard ESCRT-IIIB and ESCRT-IIIA were recruited to SLB membranes after simultaneous addition (1 μM each protein). Scale bar = 5 μm. G) Fluorescence-based quantification of ESCRT-IIIB and ESCRT-IIIA signals over time of the experiment shown in (E) (lines represent mean fluorescence intensity and shades are the SD; n=9 measured SLBs). H) Graph shows the dynamics of Asgard ESCRT-IIIB recruitment when added to the microscopy chamber alone (black dashed line: mean fluorescence. n=9) or together with ESCRT-IIIA (green line and shade: mean fluorescence and SD, respectively. n=7). ESCRT-IIIA recruitment kinetics is shown in magenta (line and shade: mean fluorescence and SD, respectively. n=9). In all cases, n represents independent SLBs.

We suspected that the inability of Asgard ESCRT-IIIA to bind flat membranes was due to the intrinsic preference of ESCRT-IIIA to form helical filaments, as is the case for its eukaryotic homologues.^42,44,46,60,63^ To test this idea, we visualised Asgard ESCRT-IIIA polymers in the absence of membranes by negative stain EM and found that they also formed thin, irregular helical filaments (Figure 5B). To determine if this intrinsic curvature correlates with a preferred membrane geometry, we used the same assay as before and added fluorescently labelled Asgard ESCRT-IIIA to NTs pulled from GUVs. Under these conditions, ESCRT-IIIA formed punctae along the NTs (Figure 5C). However, we were unable to detect similar fluorescence intensities on the body of the GUV (Figure 5C). Similar to CHMP2A-CHMP3,^60^ this indicates that ESCRT-IIIA has a clear binding preference for curved membranes. Taking this further, when ESCRT-IIIA was added to LUVs or nanorods and imaged by negative stain EM, tight helical filaments were seen decorating the membranes (Figure 5D). The diameter of the decorated membrane tubes were 31.06 ± 2.69 nm and 43.18 ± 2.21 nm, respectively (Figure S6A), similar to the diameter of membrane tubes decorated by A-type CHMP1B polymers (∼28 nm).^44^ Thus, while Asgard ESCRT-IIIB polymers are able to bind membranes with a wide range of curvatures, ESCRT-IIIA preferentially binds membranes with higher curvature - a common feature of A-type ESCRT-III proteins.

Since ESCRT-IIIA was unable to bind flat membranes, we next tested the hypothesis suggested by the phylogenetic analysis that ESCRT-IIIB would aid its recruitment (Figure 1C). As predicted, when SLBs were coated with preassembled ESCRT-IIIB, ESCRT-IIIA was specifically recruited to ESCRT-IIIB patches (Figure 5E). Interestingly, when the two proteins were added together onto flat SLBs, similar patches containing both proteins appeared (Figure 5F) with kinetics similar to rates of ESCRT-IIIA binding to pre-assembled ESCRT-IIIB (∼160 seconds to reach plateau) (Figure 5G). This was much faster than the rates observed for ESCRT-IIIB self-assembly alone (> 1000 s to reach plateau) (Figure 5H, S6B), indicating that the two proteins act synergistically to bind and co-polymerise on membranes. This may occur by mutual association and/or via the ability of each protein to alter the shape of the membrane in a way that favours the binding of the other.

Previous theoretical studies have suggested that the sequential binding of ESCRT-III proteins with different curvature preferences can drive concomitant changes in both membrane curvature^21^ and filament structure.^48^ Furthermore, simulations suggest that as they pass through intermediate states, these hetero-polymeric ESCRT-III composites transform flat membranes into tubes.^48,68,69^ To test if such a graded shift in structure is seen in our experimental data, we used negative stain EM to determine the impact of adding the two proteins to membranes. When Asgard ESCRT-IIIB was incubated alone with LUVs, polymer arrays were observed on the tubulated membranes that had an angle of 30-60° relative to the longitudinal membrane tube axis (Figure 6A, 6B). By contrast, when ESCRT-IIIA was added to membranes together with ESCRT-IIIB filaments were preferentially found oriented perpendicular to the tube axis (∼90°) (Figure 6A, 6B). While the average width of the membrane tubes appeared unchanged under these conditions, the distribution of tube sizes widened. This shift in orientation was also accompanied by a reduction in the average distance separating adjacent protofilaments in the arrays (2.53 ± 0.53 nm, down from 3.13 ± 0.53 nm with ESCRT-IIIB alone) (Figure 6C).

**Figure 6.**
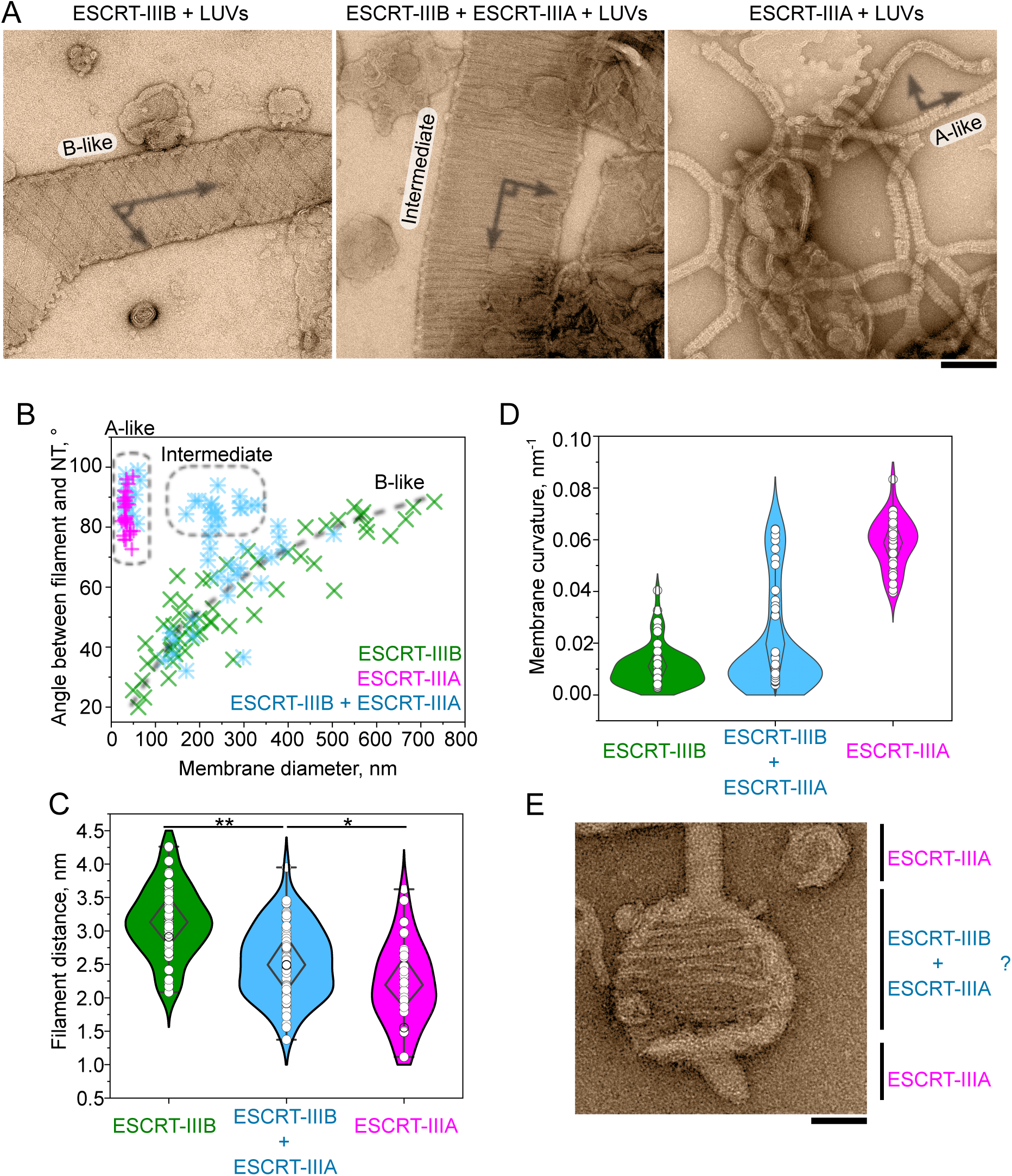
Composite polymers generated by the co-addition of ESCRT-IIIB and ESCRT-IIIA to membranes. A) Representative negative-stain EM micrographs of ESCRT-IIIB alone (left), the ESCRT-IIIB-ESCRT-IIIA co-complex (middle), and ESCRT-IIIA alone (right) upon incubation with membranes. Scale bar = 150 nm. Drawn lines over the micrographs indicate the angle taken to calculate the filament orientation shown in panel (B). B) Measurement of filament orientation defined by the angle between the membrane and filament axis (shown in A). Green and magenta crosses indicate experiments including only ESCRT-IIIB (n=57) or -IIIA (n=30) in the presence of membranes, respectively. Light blue asterisks indicate orientations measured for the ESCRT-IIIB and ESCRT-IIIA mixed sample (n=57) with membranes (n = measured filaments). Dashed boxes indicate the preferential orientations of ESCRT-IIIA alone (left) and ESCRT-IIIB-ESCRT-IIIA co-polymers (right); the dashed grey line shows the preferential orientation of ESCRT-IIIB filaments relative to the membrane diameter. C) Inter-filament distance measured from EM micrographs of ESCRT-IIIB (n=58), ESCRT-IIIA (n=43) and the mixture of ESCRT-IIIB and -IIIA (n=63) decorating membranes. (Statistical significance: unpaired two-tailed *t* test, \*\**p*<0.01; \**p*<0.05). D) Curvature estimates of the tubulated membranes decorated with ESCRT-IIIB (n=57), ESCRT-IIIA (n=30) and in ESCRT-IIIB and -IIIA mixtures (n=57) analysed from EM micrographs (n = measured filaments). E) Representative micrograph obtained by negative stain EM showing the membrane curvature in the presence of ESCRT-IIIA and ESCRT, transitioning between less curved (likely decorated with both proteins) and highly curved (probably decorated with only ESCRT-IIIA filaments) regions. Scale bar = 50 nm.

These data suggest that B-type and A-type ESCRT-III proteins can interact with one another to form composite polymers whose curvature preference allows them to occupy intermediate points along the path of membrane deformation. Interestingly, when both proteins were added simultaneously on LUVs, three populations of membrane curvature appeared (Figure 6B, 6D). One population with the lowest curvature values resembled ESCRT-IIIB alone (“B-like”). On the other hand, one population with the highest membrane curvature resembled ESCRT-IIIA alone (“A-like”). The remaining population exhibited curvatures between these two populations, indicating the existence of ESCRT-IIIB - ESCRT-IIIA co-polymers that can support “intermediate” membrane curvature. In some cases, we observed these mixed arrays transitioning into A-like tubes (Figure 6E), supporting the idea that ESCRT-IIIB containing polymers can nucleate homopolymers of ESCRT-IIIA. Taken together, these data indicate how stepwise changes in the composition of ESCRT-III polymers in the two-component system from Heimdallarchaeota progressively increase membrane curvature.

## Discussion

Since the discovery of ESCRT-III proteins in eukaryotes,^7,12^ numerous studies have examined ESCRT-III structure, biochemistry, and ESCRT-III-dependent membrane remodelling. While there were a few unique findings in these data, including polymers that contain subunits in the closed conformation,^42,44,45^ which have different membrane binding interfaces,^42,44,66^ or engage with membranes of opposite curvature to those in the cell,^49,50^ a common set of features has emerged.

Core attributes of ESCRT-III include the following: I) Most ESCRT-III polymers possess subunits in the open conformation, in which neighbouring subunits in each protofilament engage in intimate interactions with 3 to 4 subunits ahead; II) ESCRT-III protofilaments can simultaneously associate with negatively-charged lipids and with one another to generate two- and three-dimensional polymers that coat membranes; III) Flexibility of the individual ESCRT-III monomers can dictate how protofilaments adopt to distinct curvatures; IV) Different ESCRT-III proteins can function as composite polymers and heteropolymers, inducing different spontaneous curvatures to help remodel flat membranes into tubes.^49^

Here, through our analysis of a simple two-subunit ESCRT-III system identified in the genome of one of our closest prokaryotic relatives, Heimdallarchaea, we demonstrate that eukaryotes inherited all these structural features from their archaeal ancestors. Our phylogenetic analysis also shows that the entire set of eukaryotic ESCRT-III proteins can be organised into two families, which we term the B-type (before) and A-type (after) proteins. This organisation can be found in the two subunits encoded by the genomes of many Asgard archaea, ESCRT-IIIB and ESCRT-IIIA, which act in sequence to transform a flat membrane into a tube, an activity that in eukaryotes requires a much larger set of ESCRT-III proteins.

### ESCRT-III protofilament structure and binding to membranes

Our structural analysis of an Asgard ESCRT-IIIB polymer, both on and off membranes, constitutes the first detailed structure of a full-length B-type polymer (Figure 7), since previous B-type structural models only included incomplete protein chains.^43,51,52^ The different subunit conformations that we observe in our two ESCRT-IIIB structures suggest that intrinsic flexibility enables polymers to adapt to different membrane geometries and curvatures.

**Figure 7.**
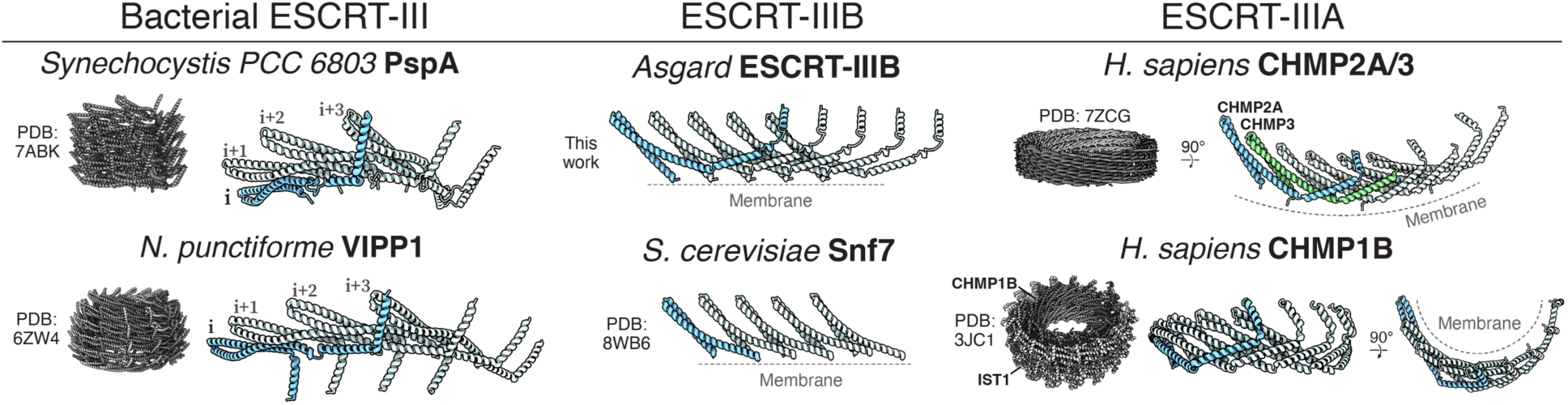
Comparison of the ESCRT-III protofilament structures across the tree of life. Figure uses structural analysis of Asgard ESCRT-IIIB to show the three main types of ESCRT-III polymers found across the tree of life: bacterial ESCRT-III; archaeal and eukaryotic ESCRT-IIIB; and archaeal and eukaryotic ESCRT-IIIA. Bacterial ESCRT-III includes *Synechocystis PCC 6803* PspA (PDB: 7ABK)^57^ and *N. punctiforme* Vipp1 (PDB: 6ZW4).^1^ ESCRT-IIIB includes our structure from Asgard and *S. cerevisiae* Snf7 (PDB: 8WB6).^52^ ESCRT-IIIA includes *H. sapiens* CHMP2A-CHMP3 (PDB: 7ZCG)^46^ and *H. sapiens* CHMP1B in complex with IST1 (PDB: 3JC1).^42^

Our structure also reveals how protofilaments interact with membranes using an interface similar to that previously described for eukaryotic B- and A-type proteins.^46,52^ Hydrophobic residues at the N-terminus of ESCRT-III proteins may dip into the lipid bilayer, suggesting that this region also contributes to membrane remodelling. We note that it is also possible for the positively charged interface of helix α1 that mediates lateral interactions between adjacent Asgard ESCRT-IIIB protofilaments to act as a second membrane binding region, similar to CHMP1B polymers.^42,44,66^ Indeed, switching between these two binding modes may help remodel membranes.^48^

### Interplay between Asgard ESCRT-III and membranes

By fluorescently labelling the two Asgard proteins, we were also able to visualise their recruitment to artificial membranes. While the precise nature of Asgard cell membranes has yet to be determined, this revealed a requirement for a critical concentration of ESCRT-IIIB protein to initiate polymer formation on flat membranes. This, we hypothesise, reflects the need to form a stable 5 subunit nucleus, likely involving i to i+4 interactions, which then extends by fast polymerisation from its free ends, as is the case for other cytoskeletal elements like actin.^70^

In cells, the local protein ESCRT-III concentration is likely not high enough for spontaneous nucleation. A barrier of this type can be overcome by the local accumulation of ESCRT components at sites rich in ubiquitinated proteins, since Asgard archaea seem to possess components equivalent to the eukaryotic multivesicular body machinery, which includes ubiquitin, ubiquitination enzymes, ESCRT-I and -II homologues.^32–34^ Although it is not clear if Asgard archaea have internal compartments, this ESCRT-III-dependent pathway may function to rid cells of excess or damaged ubiquitinated membrane proteins by producing extracellular vesicles.

In this study, we show that although Asgard ESCRT-IIIA binds to highly curved membranes when added alone, it is only recruited to flat membranes that have been pre-loaded with ESCRT-IIIB. This is similar to what is seen when combining eukaryotic B-type and A-type ESCRT-III proteins,^71^ and shows how interactions between these two proteins and the membrane function to impose order on the sequence of membrane remodelling events. In addition, our data also reveal unexpected positive feedback in the system since, when added simultaneously, the ESCRT-IIIA recruited by ESCRT-IIIB can in turn increase the rate of ESCRT-IIIB recruitment to flat membranes, leading to the rapid formation of an ESCRT-III coat. Strikingly, when imaged by negative stain EM, membranes treated with both ESCRT-III proteins were found to be of intermediate curvature - effectively transforming a flat membrane into one with a shallow buckle.^48^ Once the membrane has been deformed in this way, the disassembly of ESCRT-IIIB by Asgard Vps4^34^ likely allows ESCRT-IIIA to reach its preferred radius of curvature, constricting the membrane into a tube before scission can occur. Thus, Asgard ESCRT-IIIB and ESCRT-IIIA proteins provide an excellent simple model of multi-component ESCRT-III-dependent membrane remodelling in which B-type proteins, like CHMP4B and CHMP6, recruit A-type proteins, like CHMP2A-CHMP3 and CHMP1B-IST1 to transform a flat membrane into a tube. Our work shows that this simple mechanistic principle was preserved during eukaryogenesis, as the two ESCRT-III genes inherited from Asgard archaea underwent sequential rounds of duplication and divergence to give rise to the large set of eukaryotic ESCRT-III proteins.

Finally, this analysis of the Asgard ESCRT-III system clarifies the picture of ESCRT-III-dependent membrane remodelling across the tree of life (Figure 7). These data lead us to propose that ESCRT-III had their origin in a single protein that performed a membrane repair / remodelling function in the last common ancestor of bacteria and archaea.^1^ This was then modified with time – leading to the evolution of PspA/Vipp1 in bacteria and a multi-subunit ESCRT-III in archaea. In the common ancestor of Asgard archaea and eukaryotes, this was further elaborated through the evolution of distinct B-type and A-type ESCRT-III polymers that act in sequence. Thus, a two-protein, multistep membrane remodelling process by ESCRT-III was already established in Asgard archaea, prior to mitochondrial endosymbiosis and the origin of eukaryotes. This two component ESCRT-III system was then inherited by the ancestor of eukaryotes and elaborated by gene duplication, divergence and specialisation, giving rise to the complex system present across eukaryotes today. This complexification may have emerged as a simple consequence of the expansion in the size of the eukaryotic genome.^72^ At the same time, the increase in the number of ESCRT-III subunits enabled functional specialisation of the machinery so that it was able to act at different sites in the complex eukaryotic cell, while obeying the same fundamental mechanistic rules that have been conserved over > 2 billion years of evolution.

## Methods

### Phylogenetic analysis

Eukaryotic and Asgard ESCRT-III sequences were identified by HMMER^73^ and Blastp^74^ searches and retrieved from Uniprot^75^, Genbank^76^ and Interpro^77^ databases, with the goal of collecting a broad distribution of ESCRT-III homologues. 266 eukaryotic (33 organisms) and 81 Asgard (39 species) protein sequences were used for this analysis. Sequences were aligned with MAFFT v7.05^78^ with the (L-INS-i) setting. Maximum likelihood trees were inferred using IQTREE2 (2.2.6)^79^. Model fitting was performed in IQTREE2 for the focal analyses, with the inclusion of additional complex models (LG+C10-C60+F+G)^80^ to the default list of models. We included 10,000 ultrafast bootstraps replicates^81^, and optimised weights of mixture models. For the topology tests, constraint trees were built using TreeViewer^82^ and constrained topologies were inferred using the LG+C20+F+G model in IQTREE2. These constrained topologies were then compared using the AU test (p = 0.05).^83^

### Gene cloning, protein expression, purification and chemical labelling

Genes coding for Asgard ESCRT-IIIA and ESCRT-IIIB (Uniprot accessions A0A1Q9PC75 and A0A1Q9PC98, respectively) were cloned into the pET-28a vector (Novagen) with a His-tag and a plant SUMO domain (bdSUMO from *Brachypodium distachyon*) as N-terminal fusions.^84,85^ Plasmids were used to transform *E. coli* Rosetta DE3(pLysS) (Novagen) for recombinant protein expression. Cells were grown in 2xTY at 37°C and protein expression was induced for 4 h by the addition at mid log phase of 0.5 mM IPTG (isopropyl-β-D-thiogalactoside). Cells were harvested by centrifugation and lysed by sonication, the cleared lysate loaded onto a Histrap HP 1 mL affinity column (Cytiva) using buffer A (20 mM Tris-HCl pH 8.0, 500 mM NaCl, 5% (v/v) glycerol) and proteins eluted by a linear gradient of buffer A containing 500 mM imidazole-HCl pH 8.0. The N-terminal His-bdSUMO was removed by the *Brachypodium distachyon* SUMO protease (bdSENP1).^84,85^ After tag removal, the resulting polypeptides are the full-length versions of the Asgard proteins without any extra residues. The untagged samples were further purified using a Hiload 16/600 Superdex 200 size exclusion column (Cytiva) equilibrated with buffer A. For the fluorescence microscopy experiments, purified proteins were chemically labelled at the N-terminus with TFP-AlexaFluor-488 and/or AlexaFluor-568 NHS ester (ThermoFisher Scientific) following the procedure provided by the manufacturer.

### Analytical size exclusion chromatography

A Hiload 16/600 Superdex 200 size exclusion column (Cytiva) equilibrated with buffer A (20 mM Tris-HCl pH 8.0, 500 mM NaCl, 5% (v/v) glycerol) was used for analytical size exclusion experiments. The calibration curve is presented in Figure S2B, and was established using the following standard proteins (Merck MWGF1000): carbonic anhydrase (CAN; 29 kDa), bovine serum albumin (BSA; 66 kDa), alcohol dehydrogenase (ADH; 150 kDa), beta-amylase (BAM; 200 kDa), apoferritin (AFE; 443 kDa) and thyroglobulin (TGL; 669 kDa).

### LUV membrane and galactocerebroside nanorod preparation and generation of filaments on and off membranes

LUVs were prepared by mixing 1,2-dioleoyl-sn-glycero-3-phosphocholine (DOPC) and the negatively charged lipid 1,2-dioleoyl-sn-glycero-3-phospho-L-serine (DOPS) (Avanti) in chloroform in a molar ratio of 6:4, followed by evaporation and resuspension in buffer A (20 mM Tris-HCl pH 8.0, 500 mM NaCl, 5% (v/v) glycerol) to a final concentration of 1-2 mM. Lipids were then subjected to 10 cycles of vortexing, freezing in liquid nitrogen and thawing. Nanotubes were prepared following a similar protocol, but using a molar ratio of 4 DOPS : 2 DOPC : 4 D-Galactosyl-β1-1′-N-Nervonoyl-D-erythro-sphingosine (galactosyl ceramide; Avanti) and without the freezing and thawing cycles. Membranes were aliquoted, flash frozen and kept at -70°C until use.

To generate Asgard ESCRT-III filaments, proteins at 20 µM in buffer A (20 mM Tris-HCl pH 8.0, 500 mM NaCl, 5% (v/v) glycerol) were dialysed overnight at 4°C in the absence or presence of 200 – 2000 µM of lipids forming LUVs or NTs. The dialysis buffer was 20 mM Bis-Tris-HCl pH 6.0 for Asgard ESCRT-IIIB and 20 mM HEPES-NaOH pH 7.0 or 20 mM Tris-HCl pH 8.0 in the case of Asgard ESCRT-IIIA with or without membranes, respectively.

### Negative staining electron microscopy

Negative staining grids were prepared using a standard protocol,^86^ pipetting 3-8 µL samples on glow discharged ultrathin carbon film supported copper grids (EMS CF400-Cu-UL), followed by staining with 2% (w/v) uranyl acetate for 1 minute. Data collections were carried out in FEI Tecnai G2 Spirit and Thermo Fisher Talos L120C LaB6 cathode TEM microscopes operated at 120 Ke*V,* equipped with Ultrascan 1000 (Gatan) and CETA (Thermo Fisher) detectors, respectively.

### Cryo-EM sample preparation and data collection

Asgard ESCRT-IIIB filaments generated in the presence of 6 DOPC : 4 DOPS LUVs were applied onto 300-mesh Quantifoil R1.2/1.3 holey carbon gold grids, which were glow discharged for 12 seconds at 30 mA in an Edwards sputter coater S150B. Aliquots of 2.5 µL were pipetted on the glow discharged grids, blotted with filter paper for 3-7 seconds and force -15 N, and plunge-frozen in liquid ethane using a Vitrobot Mark IV (Thermo Fisher) at 20°C and 95% humidity. Grids were screened using a Glacios 200 kV microscope equipped with a Falcon III detector (Thermo). Cryo-EM movies were acquired at 105,000X magnification (0.826 Å/pixel) on a Titan Krios G3 300 kV microscope (Thermo), equipped with an X-FEG, 100 µm objective aperture, Gatan BioQuantum energy-filter (20 eV slit width), and K3 direct electron detector (Gatan) operated in counting mode. 29600 movies (50 frames, 1 e^-^/Å^2^/frame) at a defocus range of -0.6 to -2.6 µm were recorded using automated data collection with EPU (Thermo Fisher). The same dataset was used to solve the structure of Asgard ESCRT-IIIB bundles and membrane-bound arrays.

### Cryo-EM image processing of ESCRT-IIIB filaments

Movies were drift-corrected using MotionCorr2^87^ implemented in Relion 4.0^88^ to generate micrographs. The contrast transfer function (CTF) parameters were estimated using CTFFIND4.^89^ These micrographs were used to solve the structure of both membrane unbound bundles and membrane bound arrays.

Bundles were picked from the micrographs using the filament option in crYOLO 1.7.5 ^90^ with a model that was trained on manually picked micrographs (∼100). Manual inspection of the micrographs revealed a 31 Å repeating pattern, which was used to evenly space the picks along the bundles. ∼4.4 million particle images were extracted from the coordinates at 2.5 Å/pixel (100 pixel box size) in Relion. 2D classification with 120 classes, “ignore CTF until first peak”, and shifts restricted to 31 Å along the helical axis revealed well defined classes representing ∼760,000 particles. These were used to generate a 3D model *de novo* using InitialModel in Relion with 4 classes and imposing helical restrictions as additional arguments (--helix --helix_outer_diameter 130 --ignore_helical_symmetry --helical_keep_tilt_prior_fixed). The map that clearly resembled the 2D class averages was used to perform a symmetry search with relion_helix_toolbox,^91^ which revealed a helical symmetry of 15.5 Å rise and -178.5° twist. A less stringent selection of the 2D classes provided ∼4.1 million particles, which were used to refine against the InitialModel map.

The particles and map were imported into CryoSPARC 4.2.1^92^ for a Helix Refine job with the “align to symmetry axis” option and dynamic mask, resulting in a 5.2 Å resolution map. Particles were then converted back to Relion using csparc2star.py from pyem^93^ and extracted to 1.8 Å/pixel (188 pixel box size) with recentering according to the refined angles and shifts. A 3D Refinement in Relion using a mask covering 55% of the Z axis, where helical symmetry parameters were searched and imposed, resulted in a 3.7 Å resolution map. To isolate high-quality particles, a 3D classification without alignment with 5 classes (T=50) revealed a good class with ∼442,000 particles. A 3D Refinement of this subset resulted in 3.5 Å resolution. Per-particle defocus and per-micrograph astigmatism were refined, after which processing was transferred to Relion 5.0.^94^ Another 3D Refinement with Blush regularisation^95^ resulted in 2.9 Å resolution. Bayesian Polishing was subsequently performed using trained parameters,^96^ wherein polished particles were re-extracted at 0.826 Å/pixel (500 pixel box size). 3D Refinement with a mask covering 40% of the Z-axis resulted in 2.9 Å resolution. Another round of defocus and astigmatism refinement and 3D refinement did not significantly improve the map. Therefore, a 25% Z-mask was used for signal subtraction (250 pixel box size), from which a 3D classification without alignment (8 classes, T=4) revealed a slightly better class with ∼275k particles, which were reverted back to the original box size. 3D refinement (40% mask), defocus and astigmatism refinement, and another round of 3D refinement resulted in the final map at 2.9 Å resolution (15.4 Å rise, -178.5° twist). The map was sharpened with a B-factor of -57 Å^2^.

### Model building and refinement of ESCRT-IIIB filaments

The sharpened map was used to build a model *de novo* using ModelAngelo.^97^ Different segments generated by ModelAngelo were manually stitched together in Coot 0.9.3^98^ to generate an asymmetric unit consisting of two monomers. Due to the relatively poor density of alpha helix α5 and the adjacent region of α1/ α2, an AlphaFold2 prediction^99^ was generated of the interaction using a local installation of ColabFold 1.2.0^100^ running MMseqs2.^101^ The region of high confidence relating to this interface was stitched into each monomer model. The monomer side-chains were then manually refined in Coot 0.9.3. For real-space refinement of the polymer, the asymmetric unit was copied to the helically related positions using the sym option in Chimera 1.16.^102^ Refinement of the polymer was performed in Phenix 1.20^103^ by defining two non-crystallographic symmetry groups relating to each of the two monomers in the asymmetric unit. After refinement, symmetry-related copies were removed, and poorly defined side-chains were trimmed before model validation. Problematic side-chains were manually fixed in Coot before another round of polymer refinement and monomer validation. Map to model FSC was calculated on the polymer. Colouring of the map and model, including residues that interact, was done in ChimeraX 1.7.^104^

### Cryo-EM image processing and model building of membrane-bound arrays

Motion-corrected micrographs containing membrane-bound ESCRT-IIIB arrays were picked using a crYOLO^90^ model trained against membrane edges. An inspection of the micrographs revealed a 30 Å repeating spiked pattern. Therefore, coordinates that are spaced by 30 Å were picked along the membrane edges. An extraction of particle images followed by 2D classification revealed that the membrane signal obscured the alignment of the protofilament arrays. To overcome this, the coordinates were resampled along the membrane edge to 20 Å such that they are out of phase with the 30 Å repeat. Therefore, a rolling 2D average of the particles along any given membrane edge only enhanced the diffuse membrane signal and averaged out the protein signal. This rolling average was subtracted from the micrographs, leaving the arrays intact. Coordinate resampling and micrograph subtraction were performed using scripts originally implemented for microtubule-based processing.^64^ The original coordinates could then be extracted at 2.8 Å/pixel (90 pixel box size) in CryoSPARC 4.2.1,^92^ and a 2D classification with 50 classes while restricting shifts along the helical axis resulted in classes where the arrays wrapping around the membrane edge were nicely resolved (∼579,000 particles). Another round of 2D classification resulted in a subset of ∼378,000 particles. These were used for *ab initio* reconstruction using 1 class. The resulting map and particles underwent Homogeneous Refinement (dynamic mask) followed by Local Refinement of the central five protofilaments, resulting in a map at 5.8 Å resolution, although map connectivity remained poor at this stage.

Refined particle alignments from CryoSPARC were then converted to Relion using csparc2star.py and extracted to 2.8 Å/pixel (90 pixel box size) with re-centering. These underwent 3D Refinement in Relion 5.0,^94^ resulting in a 5.6 Å resolution map. To improve the definition of the protofilaments, the particles were re-extracted away from the very edge of the membrane (1.5 Å/pixel, 192 pixel box size). The central three protofilaments were then refined using Blush regularization^95^ and helical symmetry (also used for all subsequent refinements). In this case, symmetry relates the adjacent protofilaments, which stack laterally in the Z axis (30.1 Å rise, 0.1° twist). The resolution at this stage was 6.5 Å, with improved connectivity between alpha-helical densities. A 3D classification without alignment with 4 classes (T=4) was then performed, which revealed a good class with ∼33,000 particles. These were refined and re-extracted at 1.5 Å/pixel (274 pixel box size), shifting slightly further from the membrane edge again. To remove picks that might relate to the same particle (i.e. those that the software may have inadvertently picked twice on the same membrane edge), coordinates that were within 15 Å of each other were removed. A final 3D refinement of the central four protofilaments resulted in 6.5 Å resolution (29.9 Å rise, 0.4° twist). The map was sharpened using a B-factor of -80 Å^2^.

To build the membrane-bound array model, we used our high-resolution model of the ESCRT-IIIB bundle. One protofilament was extracted from the model and flexibly fit into the central protofilament of the membrane-bound array map in Coot 0.9.3^98^ using secondary structure restraints. Real-space refinement was performed in Phenix 1.20 before the backbone was trimmed in regions where the map was poorly resolved. To identify the interacting faces, copies of the central protofilament model were generated using the sym command in Chimera 1.16.^102^ Map to model FSC was calculated on the model that has adjacent protofilament copies. Map and model colouring, including hydrophobicity and electrostatic potential, was performed in ChimeraX 1.7.^104^

### Lipid-covered silica beads preparation

SLBs, GUVs and membrane NTs pulled from GUVs were prepared from lipid lamellae deposited on 40 mm silica beads, as previously described.^105^ 1,2-dioleoyl-sn-glycero-3-phosphocholine (DOPC), the negatively charged lipid 1,2-dioleoyl-sn-glycero-3-phospho-L-serine (DOPS) and (1,2-Dioleoyl-sn-glycero-3-phosphoethanolamine labelled with Atto 647N) Atto-647DOPE (Avanti Lipids) were mixed at either 59.95 : 40 : 0.05 or 79.95 : 20 : 0.05 mol% ratios from lipid stocks purchased from Avanti dissolved in chloroform to a final concentration of 1 mg/mL. The lipid mixture was then dried in vacuum for no less than 2 hours allowing complete chloroform evaporation forming a dried lipid film. The lipid film was hydrated and resuspended in 25 mM HEPES-NaOH buffer at pH 7.4 forming multilamellar vesicles (MLVs). Subsequently, 10 mL of the suspension was mixed with 1 mL of 40 mm silica beads (Microspheres-Nanospheres), divided into 5 drops and placed on a clean parafilm surface. Beads-MLVs drops were then dried in vacuum for at least 1 hour until complete evaporation of the buffer.

### SLB membrane preparation and TIRF microscopy on Asgard ESCRT-III protein recruitment

SLBs were prepared as previously described.^105,106^ Initially, a coverslip was cleaned in water/ethanol/water for 10 minutes each using sonication. Following this washing step, coverslips were completely dried using nitrogen gas and plasma cleaned (Harrick Plasma) for 30 seconds. An *Ibidi* chamber (sticky-Slide VI 0.4) was then mounted on the coverslip and filled up with working buffer (25 mM HEPES-NaOH, 10 mM MgCl_2_ at pH 6.5). Lipid-covered silica beads (See *Lipid-covered silica beads preparation* method section) were transferred using a 20 mL plastic micropipette tip to the wells used for the experiments. After 10-15 min incubation, lipid bilayers were spilled on the coverslip, leading to the formation of the SLBs. Fluorescently labelled proteins (at concentrations and combinations specified in each experiment) were added to the *Ibidi’*s inlet using a syringe pump withdrawing the bulk volume from the outlet position of the flow chamber. Sequential addition of ESCRT-IIIB and -IIIA experiment was performed by first incubating ESCRT-IIIB with the SLBs until the formation of ESCRT-IIIB patches, then washing-out bulk ESCRT-IIIB to stop its polymerization before complete membrane surface coverage, and finally adding ESCRT-IIIA in the incubation chamber. Simultaneous ESCRT-IIIB and ESCRT-IIIA addition was performed by adding both proteins at the same time in the incubation chamber.

TIRF microscopy experiments were performed using an Olympus IX83 wide-field microscope equipped with an Olympus Uapo N 100X 1.49 oil objective and an ImageEM X2 EM-CCD camera (Hamamatsu). The system was controlled by the Visiview v4.4.0.11 software (Visitron Systems GmbH).

### Quantification of the rate of fluorescence increase in the polymerisation regime and the maximum surface coverage by Asgard ESCRT-IIIB

The fluorescence increase rate for polymerisation was obtained from the slope of the linear region of the fluorescence plot profile (Figure 3B) at the concentrations at which ESCRT-IIIB polymerises (300, 500, 750 and 1000 nM). Maximum surface coverage by ESCRT-IIIB was obtained by fitting the experimental data from Figure 3C with a sigmoid function and extracting the value corresponding to the maximum fluorescence intensity at the steady-state.

### Preparation and visualisation of GUV and GUV-pulled lipid NTs incubated with Asgard ESCRT-III proteins

GUVs were prepared using lipid-covered silica beads (See *Lipid-covered silica beads* preparation method section), as described earlier.^105^ Lipid-covered silica beads were hydrated in a 1 M trehalose solution for 15 min at 60°C in a home-made humidity chamber and deposited in the microscopy observation chamber. At this stage, growing GUVs were still attached to the silica beads. Therefore, in order to generate free-standing GUVs, the chamber was gently stirred manually for 1 min promoting the detachment of GUVs from the silica support beads. Fluorescently-labelled protein (concentration and subunit composition specified on each particular experiment) was carefully added after free-standing GUV production. For pulling membrane NTs from free-standing GUVs, closed glass micropipettes were initially prepared using a P-1000 micropipette puller (Sutter Instruments, USA). Lipid NTs were produced by direct physical contact between the micropipette and the GUVs. Micropipettes were moved in XYZ inside the microscopy chamber using a micro-positioning system (MP-285, Sutter Instrument, Novato, CA, USA). Fluorescently-labelled protein was added immediately before tube pulling to the microscopy chamber (concentration indicated in each experiment). In both cases, fluorescence image acquisition was carried out using an inverted spinning disc microscope assembled by 3i (Intelligent Imaging Innovation), composed by a Nikon base (Eclipse C1, Nikon), a 100x 1.49 NA oil immersion objective, and an EVOLVE EM-CCD camera (Roper Scientific).

### Measurement of Asgard ESCRT-IIIB relative abundance between curved and flat membranes

Relative abundance of fluorescently labelled ESCRT-IIIB was obtained by extracting the plot profile of the protein signal in the curved and flat regions of the membrane (GUV and pulled lipid nanotube, respectively). Plot profile data-points were then normalised against plot profiles of the fluorescence intensity obtained from the membrane at the exact same ROIs as previously used to measure ESCRT-IIIB fluorescence intensity.

### Quantification of ESCRT-III-induced membrane structural features

Quantification of Asgard ESCRT-IIIA filament thickness after incubation with LUVs and nanorods was performed by measuring the maximum distance between the outer edges of the protein filaments. The relative orientation of the distinct ESCRT-III filaments (either ESCRT-IIIB alone, ESCRT-IIIA alone, or the mixture of both) was obtained by measuring the smaller angle between the membrane longitudinal axis and filament decorating the membrane (as indicated in Figure 6A by the arrows). Filament distances from samples with different Asgard ESCRT-III subunit composition were obtained by measuring the peak distances in their plot profiles perpendicular to the filament longitudinal axis.

## Acknowledgments

We would like to thank the staff of the MRC LMB and DCI Geneva EM facilities, Jan Löwe and Sjors Scheres for support on electron microscopy sample preparation, data collection and cryo-EM image processing. We would also like to thank Fréderic Humbert and Rafael Ferreira in the Roux lab for technical support on protein expression and purification, Andrew P. Carter for insightful discussions and suggestions, and Jake Grimmet and the Scientific Computing team at the MRC LMB for providing computing resources. We would like to thank the following people for their critical feedback on the manuscript: Kris Kuo, Jan Löwe, Adam Prada, Andela Saric and Josh Tran. This work was funded by Wellcome Trust funding to BB, which supported DPS (203276/Z/16/Z), and via MRC support for the Baum team. JE is supported by a long-term EMBO fellowship (EMBO ALTF 989-2022). AR acknowledges funding from the Swiss National Fund for Research grant numbers #CRSII5_189996 and #310030_200793 and the European Research Council Synergy grant number #951324-R2-TENSION. SC is supported by a Wellcome Trust grant to Andrew P. Carter (210711/Z/18/Z). TH was supported by a Wellcome Trust collaborative award jointly held by MB and BB (203276/Z/16/Z). TW and EM were supported by a grant from the Gordon and Betty Moore Foundation, GBMF9741.

## Author contributions

DPS collected protein sequences for phylogenetic studies, proposed the evolutionary history of Asgard and eukaryotic ESCRT-III, purified proteins, performed the analytical SEC analysis, obtained and optimised the ESCRT-IIIA and -B filaments and prepared Cryo-EM samples. JE carried out fluorescence microscope experiments on SLBs, pulled lipid NTs and GUVs, and all associated measurements and interpretations. JE and DPS carried out negative stain electron microscopy assays. DPS and SC collected the cryo-EM data and interpreted structures. SC performed cryo-EM image processing and model building, with support and input from DPS. ERRM and TAW generated the phylogenetic tree and DPS, ERRM and TAW interpreted them. DPS and TH performed gene cloning. TH initiated biochemistry and studies of ESCRT-III polymerisation. MB supervised TH. DPS, JE, AR and BB designed the study. DPS, JE and SC generated the figures. DPS, JE, SC, AR and BB wrote the paper, with input from all authors. BB and AR supervised the work and obtained funding. The authors wish it to be known that, in their opinion, the three first authors (DPS, JE and SC) should be considered as joint first authors and as having made equal contributions to this work.

## Data availability

Atomic coordinates and cryo-EM maps have been deposited in the Protein Data Bank (PDB) / Electron Microscopy Data Bank (EMDB) under accession codes 9FTL/50748 (off-membrane four-protofilament bundle) and 9FTM/50749 (on-membrane array).

