## Supplementary Figures for "Evolutionarily conserved principles of ESCRT-III-mediated membrane remodelling revealed by a two-subunit Asgard archaeal system"

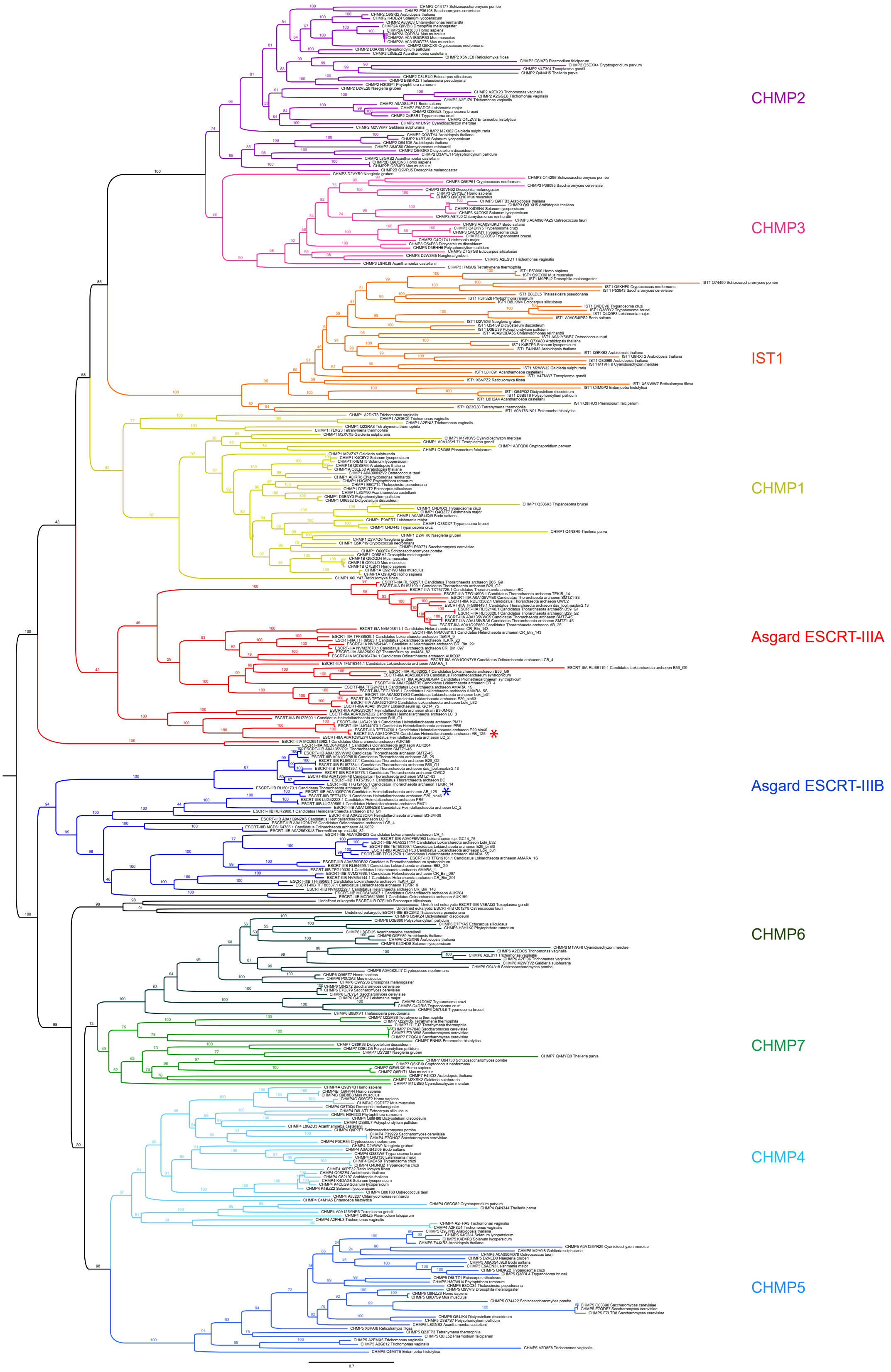


**Figure S1**. Complete phylogeny of the eukaryotic and Asgard ESCRT-III subfamilies, related to Figure 1. This is the same tree shown in Figure 1A and using an equivalent colour scheme, showing more details for each branch. For each sequence, the corresponding ESCRT-III subfamily, its UniProt or GenBank code, and the species in which it is found are described. Numbers show the relative support from 10,000 bootstrap replicates. Scale bar represents expected substitutions per site. The A- and B-type sequences from Heimdallarchaeota archaeon AB_125, investigated experimentally in this study, are labelled with red and blue asterisks, respectively.

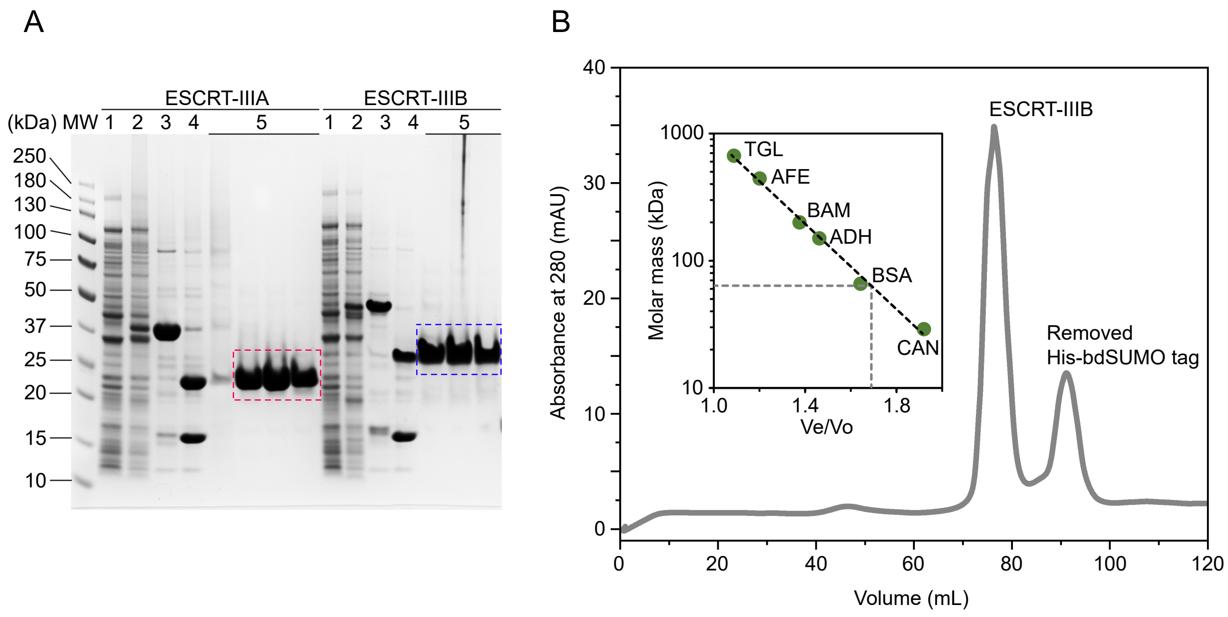


**Figure S2**. Protein purification and analytical size exclusion chromatography, related to Figures 2-6. A) SDS-PAGE of the purification of Asgard ESCRT-IIIA (left) and ESCRT-IIIB (right). MW: molecular weight (in kDa) marker; 1: and 2: *E. coli* cell extract before and after the addition of the inducer, respectively; 3: affinity chromatography fractions; 4: samples after the addition of the bdSUMO protease and tag cleavage; 5: size-exclusion chromatography fractions. Purified ESCRT-IIIA and ESCRT-IIIB are indicated by the red and blue dashed boxes, respectively. B) Superdex 200 16/600 size-exclusion profile of Asgard ESCRT-IIIB (predicted mass of its monomer: 24 kDa). The peak corresponding to the His-bdSUMO tag removed by proteolysis is also indicated. The void volume (Vo) of this column corresponds to approximately 45 mL. Inset: S200 16/600 calibration curve (dashed black line) generated using 6 standard protein markers [green dots: carbonic anhydrase (CAN; 29 kDa), bovine serum albumin (BSA; 66 kDa), alcohol dehydrogenase (ADH; 150 kDa), beta-amylase (BAM; 200 kDa), apoferritin (AFE; 443 kDa) and thyroglobulin (TGL; 669 kDa)]. The dashed grey lines represent the Ve/Vo (Ve: elution volume) and the predicted molar mass (63 kDa) of ESCRT-IIIB. This experiment indicates that this protein does not predominantly form high polymeric forms during purification and is expressed and purified as a dimer or most likely as an elongated monomer.

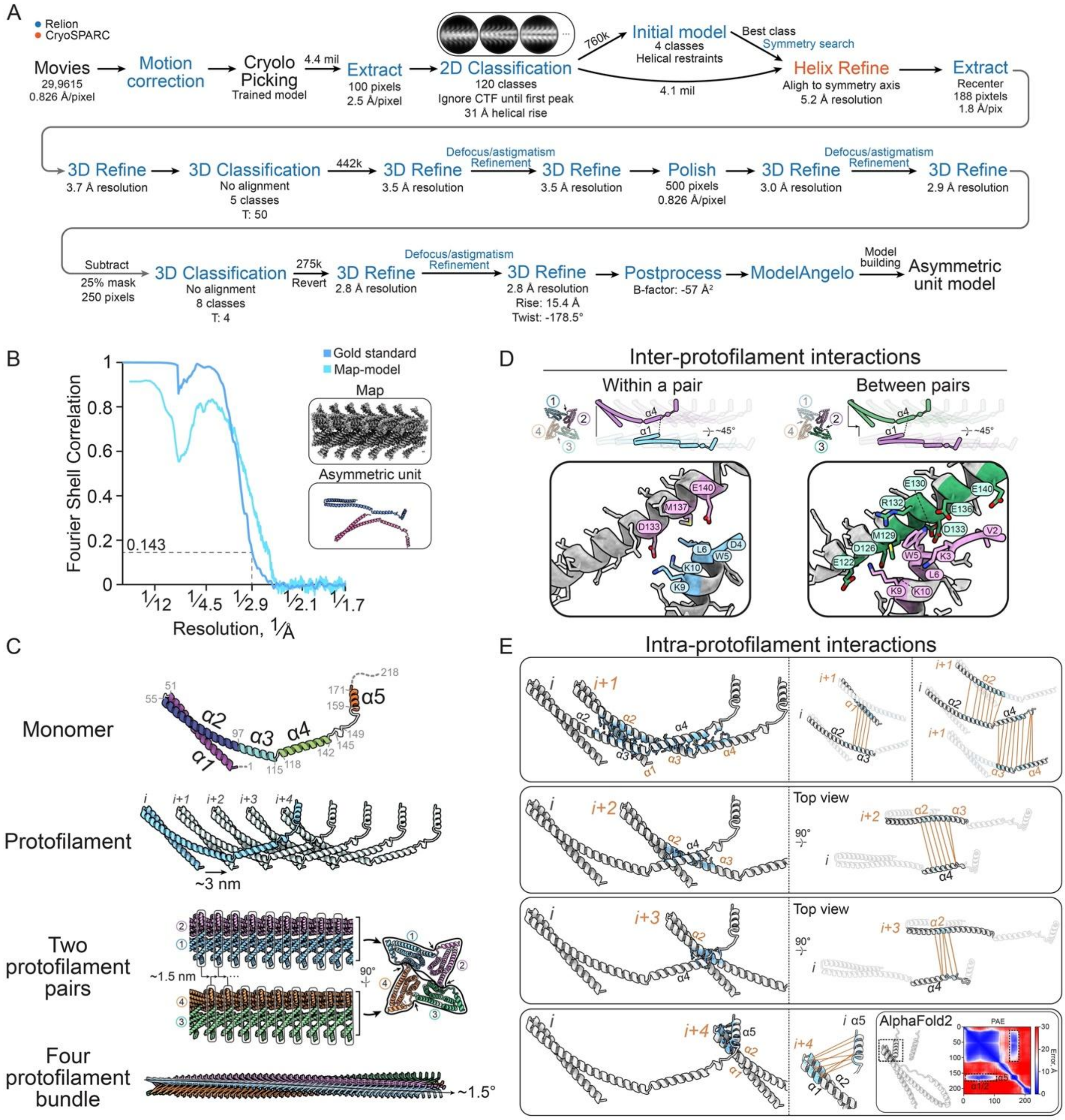

**Figure S3**. **Cryo-EM processing and structural characterisation of ESCRT-IIIB filaments, related to Figure 2.** A) Processing pipeline for single-particle analysis of Asgard ESCRT-IIIB bundles. Relion jobs are labelled in blue and CryoSPARC jobs are labelled in orange. All 3D refinements include helical refinement, where helical parameters are searched and imposed. B) Fourier shell correlation showing the 0.143 cut-off. C) Subunit architecture of Asgard ESCRT-IIIB monomers, protofilaments, and the four-protofilament bundles. Two protofilament pairs are distinguished by the different axial shift of their subunits along the helical axis. A zoomed-out view of the bundle shows the ~1.5° supertwist. D) Interaction interface between adjacent protofilaments within a pair (i.e. axially aligned) and between pairs. Note that W5 and D4 sidechains are unresolved and therefore hidden in the left-hand zoom-in. E) Overview of the interaction between adjacent subunits of a protofilament, which includes *i* to *i+1*/*i+2*/*i+3*/*i+4*. Interacting residue backbones are coloured in cyan and interfaces are separated to help visualise the interactions (orange lines). An AlphaFold2 prediction of the *i* to *i+4* interaction is shown along with its predicted aligned error (PAE) plot.

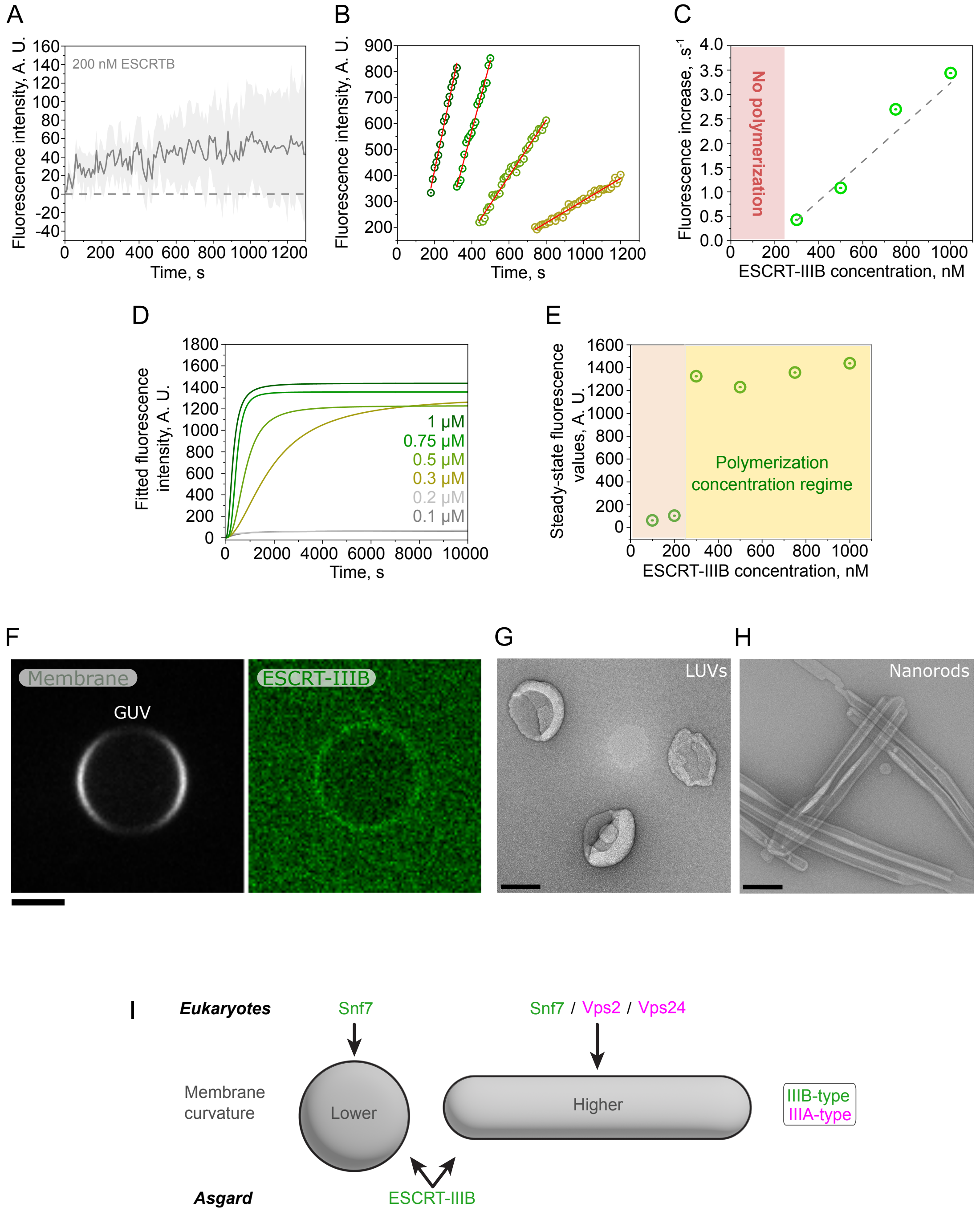


**Figure S4**. **Interaction of Asgard ESCRT-IIIB with lipid membranes. Related to Figure 3.** A) Fluorescence intensity time-profile of Asgard ESCRT-IIIB on SLBs below the polymerisation concentration regime (200 nM), lacking the characteristic sigmoidal shape of ESCRT-IIIB polymer assembly which can be seen in Figure 3B-C (line: mean; shadow: SD; n=9 measured SLBs). B) Experimental data-points and a line fit for the linear part of the polymerisation regime when ESCRT-IIIB is added to membranes at

300, 500, 750 and 1000 nM (from right to left). C) Graph shows rate of increase in fluorescence intensity versus concentration for experiments in which Asgard ESCRT-IIIB was added to membranes. Data was extracted from the linear part of the polymerisation regime (shown in Figure S4B). The grey dashed line is the line fit, indicating a linear increase of growth rate with increasing Asgard ESCRT-IIIB concentrations in this regime. D) Curves show fitted experimental data shown in Figure 3C using a sigmoidal function. E) Predicted fluorescence intensity value at steady-state, obtained from fitted curves shown in Figure S4D. Below a concentration of ~250 nM, Asgard ESCRT-IIIB fails to assemble onto membranes (light pink background). Above this concentration it polymerises on membranes until completely covering the SLB surface (yellow background). F) Fluorescence micrograph of a GUV with slight binding of ESCRT-IIIB insufficient to trigger membrane "crumpling". G-H) Representative micrographs of (G) nanorods and (H) LUVs visualised by negative stain EM. Scale bars = 120 nm. I) Schematic illustrating the ability of ESCRT-IIIB to adapt to different membrane curvatures, as compared to yeast ESCRT-III proteins where the curvature transition is promoted by the addition of Vps2/Vps24 forming a composite heteropolymer together with Snf7.

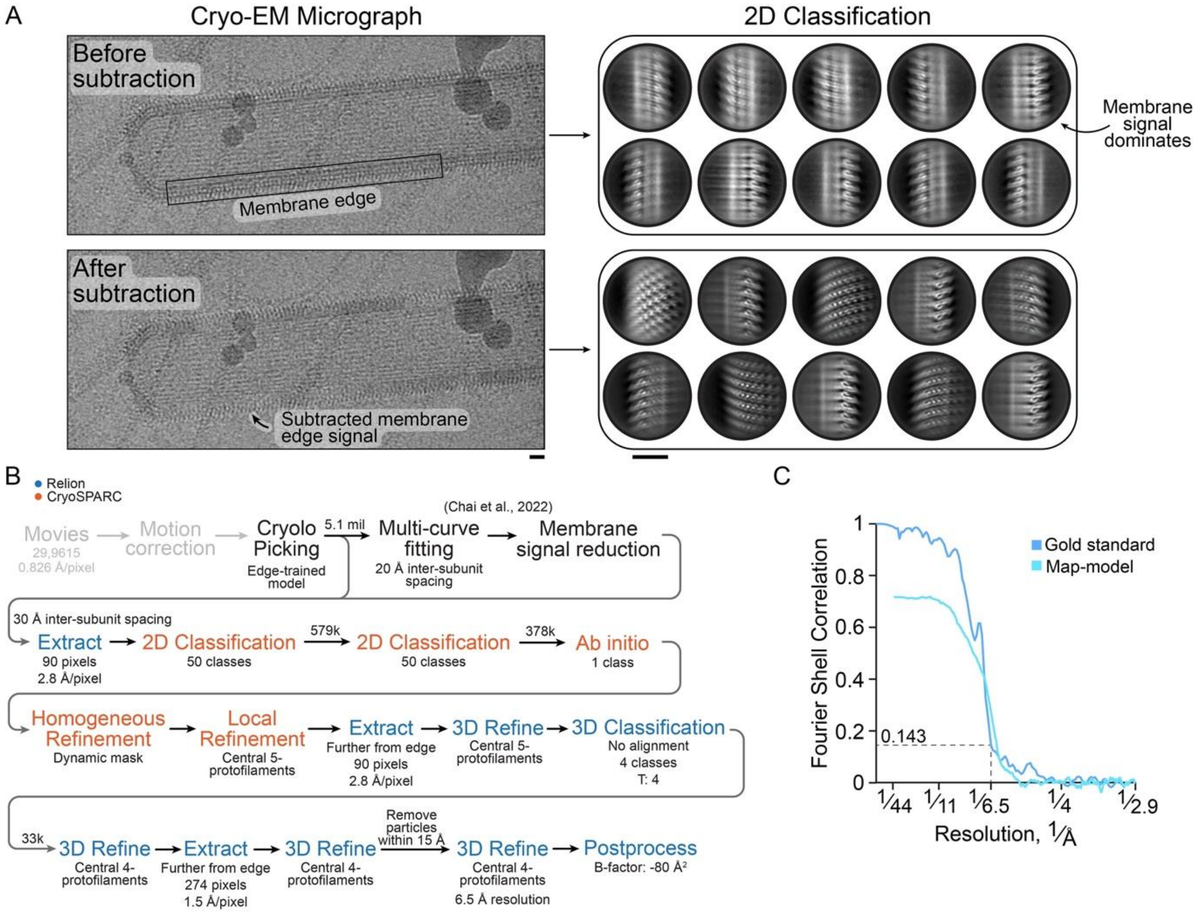


**Figure S5**. **Cryo-EM processing of ESCRT-IIIB arrays bound to membranes, related to Figure 4.** A) An example micrograph before and after subtraction of the membrane signal, as well as representative 2D class averages demonstrating the removal of the dominating signal. Scale bars are 10 nm. B) Processing pipeline for single-particle analysis of membrane-bound arrays. Relion jobs are labelled in blue and CryoSPARC jobs are labelled in orange. C) Fourier shell correlation showing the 0.143 cut-off.

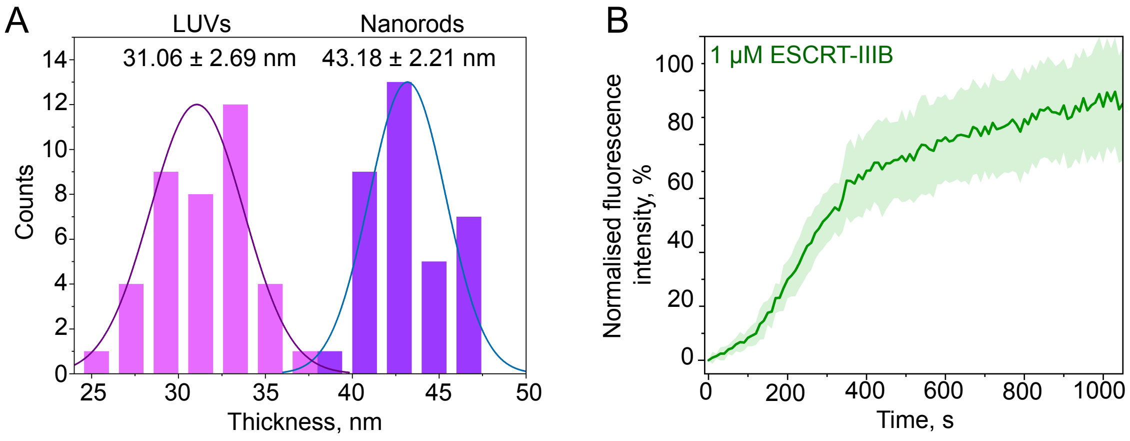


**Figure S6. Asgard ESCRT-III polymerisation on membrane surfaces. Related to Figures 5 and 6.** A) Histogram showing the thickness distribution of ESCRT-IIIA filaments polymerising on LUVs (n=39; 31.06 ± 2.69) and nanorods (n=35; 43.18 ± 2.21) (n = measured filaments; mean ± SD). B) Fluorescence profile of 1 mM Asgard ESCRT-IIIB on SLBs acquired using TIRF microscopy and normalised to its maximum fluorescence intensity. Line: mean; Shadow: SD (n=9 independent SLBs).

| | Four-protofilament bundle | Membrane-bound array |
|---|---|---|
| **PDB** | 9FTL | 9FTM |
| **EMDB** | 50748 | 50749 |
| **Data collection and processing** | | |
| Magnification | 105,000 | 105,000 |
| Voltage (kV) | 300 | 300 |
| Electron exposure ($e^{-}/Å^2$) | 50 | 50 |
| Defocus range (μm) | 1.2 – 2.6 | 1.2 – 2.6 |
| Pixel size (Å) | 0.826 | 0.826 |
| Symmetry imposed | Helical | Helical |
| Rise (Å) | 15.4 | 29.9 |
| Twist (°) | -178.5 | 0.4 |
| Initial particle images (no.) | 4.4 million | 5.1 million |
| Final particle images (no.) | 274,671 | 27,747 |
| Map resolution (Å) | 2.9 | 6.5 |
| FSC threshold | 0.143 | 0.143 |
| **Refinement** | | |
| Model resolution (Å) | 3.1 | 7.8 |
| FSC threshold | 0.5 | 0.5 |
| Map sharpening *B* factor ($Å^2$) | -57 | -80 |
| Model composition | | |
| Non-hydrogen atoms | 2,404 | 2,621 |
| Protein residues | 335 | 527 |
| *B* factors ($Å^2$) | | |
| Protein | 2404/0 | 2,621/0 |
| R.m.s. deviations | | |
| Bond lengths (Å) | 0.005 (0) | 0.004 (0) |
| Bond angles (°) | 1.083 (0) | 0.946 (0) |
| Validation | | |
| MolProbity score | 0.84 | 0.8 |
| Clashscore | 1.07 | 1.03 |
| Poor rotamers (%) | 0.45 | 0 |
| Ramachandran plot | | |
| Favored (%) | 97.89 | 98.37 |
| Allowed (%) | 2.11 | 1.63 |
| Disallowed (%) | 0 | 0 |

**Table S1. Cryo-EM data collection, refinement, and validation statistics of the four-protofilament bundle and membrane bound array.**
